# Population structure limits inferences from genomic prediction and genome-wide association studies in a forest tree

**DOI:** 10.1101/2024.10.11.617670

**Authors:** Gancho T. Slavov, David Macaya-Sanz, Stephen P. DiFazio, Glenn T. Howe

## Abstract

There is overwhelming evidence that forest trees are locally adapted to climate. Thus, genecological models based on population phenotypes have been used to measure local adaptation, assess risks of genetic maladaptation to climate, and guide assisted migration. However, instead of phenotypes, there is increasing interest in using genomic data for gene resource management. We used whole-genome resequencing and a replicated common-garden experiment to understand the genetic architecture of adaptive traits in black cottonwood. We studied the potential of using genome-wide association studies (GWAS) and genomic prediction to detect causal loci, identify climate-adapted phenotypes, and practice assisted migration. We analyzed hierarchical population structure by partitioning phenotypic and genomic (SNP) variation among 840 genotypes collected from 91 stands along 16 rivers. Most phenotypic variation (60-81%) occurred at the population level and was strongly associated with climate. Population phenotypes were predicted well using genomic data (e.g., predictive ability *r* > 0.9) but almost as well using climate or geography (*r* > 0.8). In contrast, genomic prediction within populations was poor (*r* < 0.2). Similarly, we identified many GWAS associations among populations, but most appeared to be spurious based on pooled within-population analyses. Hierarchical partitioning of linkage disequilibrium and haplotype sharing suggested that within-population genomic prediction and GWAS were poor because allele frequencies of causal loci and linked markers differed among populations. Our results highlight the difficulty of using GWAS to identify causal loci when there is population structure, and the limitations of using genomic information alone to guide assisted migration.

## Introduction

Forests are key components of global biodiversity and other important ecosystem services, including fuelwood and timber production, regulation of water and air quality, carbon sequestration, climate regulation, and spiritual and recreational experiences (MILLENNIUM ECOSYSTEM ASSESSMENT 2005). However, forests are under pressure from human population growth, conversion of forests to agricultural land, commodity production, wildfire, urbanization, and climate change (CURTIS *et al*. 2018; IPCC CORE WRITING TEAM 2023). For example, forest inventories and species distribution models suggest there will be profound shifts in habitats of tree species with climate change (GARZÓN *et al*. 2019; HILL AND FIELD 2021), likely resulting in maladaptation of locally adapted populations (WANG *et al*. 2006; ST CLAIR AND HOWE 2007; AITKEN *et al*. 2008; FRANK *et al*. 2017).

Historically, population-level genetic variation has been studied by measuring phenotypes in common gardens (LANGLET 1971; MORGENSTERN 1996; ST. CLAIR *et al*. 2019). These studies established the prevalence of clinal genetic variation along climatic gradients, local adaptation, and greater genetic differentiation for putatively adaptive traits than for neutral genetic markers (i.e., *Q*_ST_ > *F*_ST_) (REHFELDT *et al*. 1999; REHFELDT *et al*. 2002; HOWE *et al*. 2003; ST CLAIR *et al*. 2005; ALBERTO *et al*. 2013). Thus, gene flow may be extensive in forest trees (HAMRICK *et al*. 1992; SLAVOV *et al*. 2004; PETIT AND HAMPE 2006) but often appears to be outweighed by diversifying selection.

Because climate is a key driver of natural selection, genecological models have been developed to understand the relationships between climate and population-level phenotypes. Growth rate and vegetative bud phenology are typically used as phenotypes because they are consistently associated with climate, genetically correlated with adaptation to cold and drought, and considered surrogates for fitness (CAMPBELL 1979; REHFELDT *et al*. 1999; WANG *et al*. 2006; WANG *et al*. 2010; LEITES *et al*. 2012). The resulting models have been used to assess the risks of genetic maladaptation from climate change (ST CLAIR AND HOWE 2007; FRANK *et al*. 2017) and guide assisted migration (GRAY *et al*. 2011; AITKEN AND WHITLOCK 2013; GRAY AND HAMANN 2013; HAMANN *et al*. 2015; AITKEN AND BEMMELS 2016). However, these genecological models also have limitations. First, multiple long-term field trials (e.g., >10; WANG *et al*. 2010) are needed to predict field performance accurately, and this is time-consuming and costly. Second, climate-based models do not necessarily account for demographic factors. Third, within-population prediction is limited by the resolution of climate models (WANG *et al*. 2016; YE *et al*. 2022). Overall, genecological models are valuable for inferring deployment areas for breeding populations but contribute little to genetic improvement within populations. Ultimately, genomic information may help overcome some of these limitations (CHEN *et al*. 2022).

There has been a long-standing interest in using genetic markers instead of phenotypes to manage forest genetic resources. Initially, these studies focused on presumably neutral markers such as allozymes (MILLAR AND WESTFALL 1992; WESTFALL AND CONKLE 1992) but interest increased dramatically with the potential for using genetic markers associated with adaptive traits. Over the past two decades, candidate markers have been identified using association analysis of functional candidate genes (i.e., potential causal loci; INGVARSSON *et al*. 2008; ECKERT *et al*. 2009; HOLLIDAY *et al*. 2010), patterns of gene expression (ROHDE *et al*. 2007; HOWE *et al*. 2015), positions relative to mapped QTL in biparental families (BROWN *et al*. 2003; HOWE *et al*. 2003), genotype-environment associations (GEA; ECKERT *et al*. 2010a; ECKERT *et al*. 2010b), and associations with phenotypes via genome-wide association studies (GWAS; EVANS *et al*. 2014; MCKOWN *et al*. 2014b; MÜLLER *et al*. 2017; MCKOWN *et al*. 2018; WANG *et al*. 2018). Despite extensive research on candidate genes, most remain to be validated as causal loci in natural or breeding populations.

GWAS and genomic prediction methods are now widely used to study the genetic architectures of complex quantitative traits: the number, locations, and allele frequencies of causal loci, plus their additive, dominance, epistatic and pleiotropic effects (TIMPSON *et al*. 2018; MACKAY AND ANHOLT 2024). Because genetic architecture is often inferred from linked markers, we use ‘genetic architecture’ to refer to the genetic characteristics of causal loci and loci in linkage disequilibrium (LD) with causal loci. The ultimate goal of GWAS is to detect causal loci, identify potential targets for genetic modification, and help predict phenotypes. Although thousands of GWAS loci have been reported in forest trees, reproducibility has been low, and few have been validated (GRATTAPAGLIA *et al*. 2018; STRAUSS *et al*. 2022). This is because locus effect sizes are small for polygenic traits, GWAS is prone to statistical biases, population structure for quantitative traits and neutral markers are often confounded, and population sample sizes are typically low (e.g., average *N* = 446; HALL *et al*. 2016).

With low heritabilities or small sample sizes (i.e., low power), GWAS is prone to the ‘winner’s curse,’ whereby effect sizes are overestimated when the same data are used to detect statistically significant loci and estimate locus effect sizes (JOSEPHS *et al*. 2017; FORDE *et al*. 2023). With small sample sizes, GWAS are also prone to the ‘Beavis effect,’ resulting from a combination of the winner’s curse and inability to distinguish individual causal loci (BEAVIS *et al*. 1994; BEAVIS 1998; JOSEPHS *et al*. 2017). Particularly in pedigree-based studies, small sample sizes result in fewer recombination events, leading to co-segregation of linked QTL, underestimation of the number of causal loci, and inflated locus effect sizes.

Clustered loci are even harder to distinguish in genomic regions with low recombination (JOSEPHS *et al*. 2017; LOTTERHOS 2019), making it challenging to identify most of the specific loci controlling a polygenic trait. Finally, when GWAS is used among populations (e.g., range-wide studies), phenotypic variation is often confounded with neutral population structure (PRICE *et al*. 2010), leading to false positive associations. Methods exist to mitigate this problem (PRICE *et al*. 2010), but they are not ‘bulletproof’ and may also increase the rate of false negatives by over-correcting for population structure. Thus, as described for GEA (LOTTERHOS 2023), among-population GWAS suffers from a ‘Catch-22’—results may be ‘riddled with false positives’ without corrections for population structure, or causal loci will be missed when corrections are used. Despite the predominance of among-population GWAS in forest trees (STRAUSS *et al*. 2022), a substantial proportion of quantitative genetic variation resides within populations (HOWE *et al*. 2003; SCOTTI *et al*. 2016). This makes within-population GWAS informative and tractable. In humans, for example, it is common to aggregate results from within-population studies across populations using meta-analysis, rather than relying on fundamentally more challenging across-population analyses (LI AND KEATING 2014; WANG *et al*. 2020; KACHURI *et al*. 2024).

One way to predict phenotypes is to use many GWAS loci in a single prediction equation (e.g., polygenic score; WANG *et al*. 2020). This is enticing, but the loci must account for a substantial proportion of the phenotypic variation to be valuable, a tall order given that few GWAS associations have been validated in forest trees (GRATTAPAGLIA *et al*. 2018; STRAUSS *et al*. 2022). Alternatively, phenotypes can be predicted using all available markers, assuming most causative loci will be assayed directly or via LD with at least one marker, an approach called genomic selection or genomic prediction (MEUWISSEN *et al*. 2001). This approach, which focuses on prediction rather than identifying causal loci, has been widely used in animal and plant breeding populations (DAETWYLER *et al*. 2013; HICKEY *et al*. 2017; GRATTAPAGLIA *et al*. 2018), humans (MAKOWSKY *et al*. 2011; CHEN *et al*. 2015; LELLO *et al*. 2018), and more rarely, natural populations of forest trees and other plants (HOLLIDAY *et al*. 2012; KOOKE *et al*. 2016). Finally, GWAS and genomic prediction can be integrated. In addition to reflecting relatedness among individuals, genetic markers can track Mendelian segregation of linked QTL, increasing the accuracy of genomic prediction. Thus, GWAS statistics and other criteria have been used to preselect markers for genomic prediction (LING *et al*. 2021). However, using GWAS statistics is tricky—if there are too many false positives, the preselected loci may perform much worse than simply using all available loci (TAN AND INGVARSSON 2022). Furthermore, results will be biased unless GWAS, model training, and prediction are conducted in independent populations (LING *et al*. 2021; TAN AND INGVARSSON 2022), which is challenging given the large sample sizes needed to detect loci for complex quantitative traits.

Black cottonwood (*Populus trichocarpa*), a fast-growing, riparian tree, is ideal for genomic studies of adaptive traits. It occurs from Baja, California to Alaska (DEBELL 1990), inhabits diverse environments, and has well developed phenotypic and genomic resources (TUSKAN *et al*. 2006; SLAVOV *et al*. 2012; GERALDES *et al*. 2013; EVANS *et al*. 2014; MCKOWN *et al*. 2014b; HOLLIDAY *et al*. 2016). Because most of the adaptive genetic variation in black cottonwood occurs at the river level (DUNLAP AND STETTLER 1996), we sampled 1100 clonal genotypes from 23 rivers, mostly from the core of the species range in western Oregon, western Washington, and southwestern British Columbia (SLAVOV *et al*. 2012). Additionally, to strengthen within-population analyses, we sampled four of these rivers more intensively. We used phenotypic measurements from three replicated field trials and > 20M single-nucleotide polymorphisms (SNPs) from whole-genome resequencing.

Then, we used SNPs from a subset of 840 clonal genotypes from 16 rivers to address four questions: (1) How does population genetic structure (i.e., the distribution of genetic variation within vs among populations) differ between SNPs and adaptive trait phenotypes? (2) How do population differences in genetic architecture influence the ability to identify or tag causal loci using GWAS and predict phenotypes within and among populations? (3) How well are phenotypes predicted from SNPs compared to climate and geographic variables? (4) What is the potential for using genomic information from natural populations for gene conservation, breeding, and assisted migration? In contrast to other studies, we combined among- population and within-population analyses to better understand the genetic architecture of adaptive traits in forest trees.

## Results

### Phenotypic variation in adaptive traits was highly structured and strongly associated with climate

Genetic variation for adaptive traits was highly structured (Fig. 1B). For example, the correlation between latitude and the first principal component for quantitative traits (QPC1) was 0.75. Furthermore, differentiation among stands (i.e., contiguous groups of trees, typically spanning up to several kilometers) was high (*Q*_ST_ = 0.42-0.68, Fig. 2). When genotypic variation for bud flush, bud set, and height was partitioned hierarchically among river, stand-within-river, and genotype-within-stand-and-river levels, more than 50% of the variation occurred at the river level (Fig. 2, first three columns). Finally, based on multivariate regression, phenotypic variation in these traits was strongly associated with climate (*SI Appendix*, Table S1). Based on random forest analyses at the stand level, variables associated with evaporative demand and moisture deficit (Eref, CMD, SHM, MSP), temperature extremes (EXT, EMT, MCMT, TD), and winter chilling (DD<0) were consistently the best predictors of quantitative traits and SNP PC scores (SPC1-SPC5; *SI Appendix*, Table S1). Analyses at the river level resulted in similar patterns (*SI Appendix*, Table S2).

**Figure 1.**
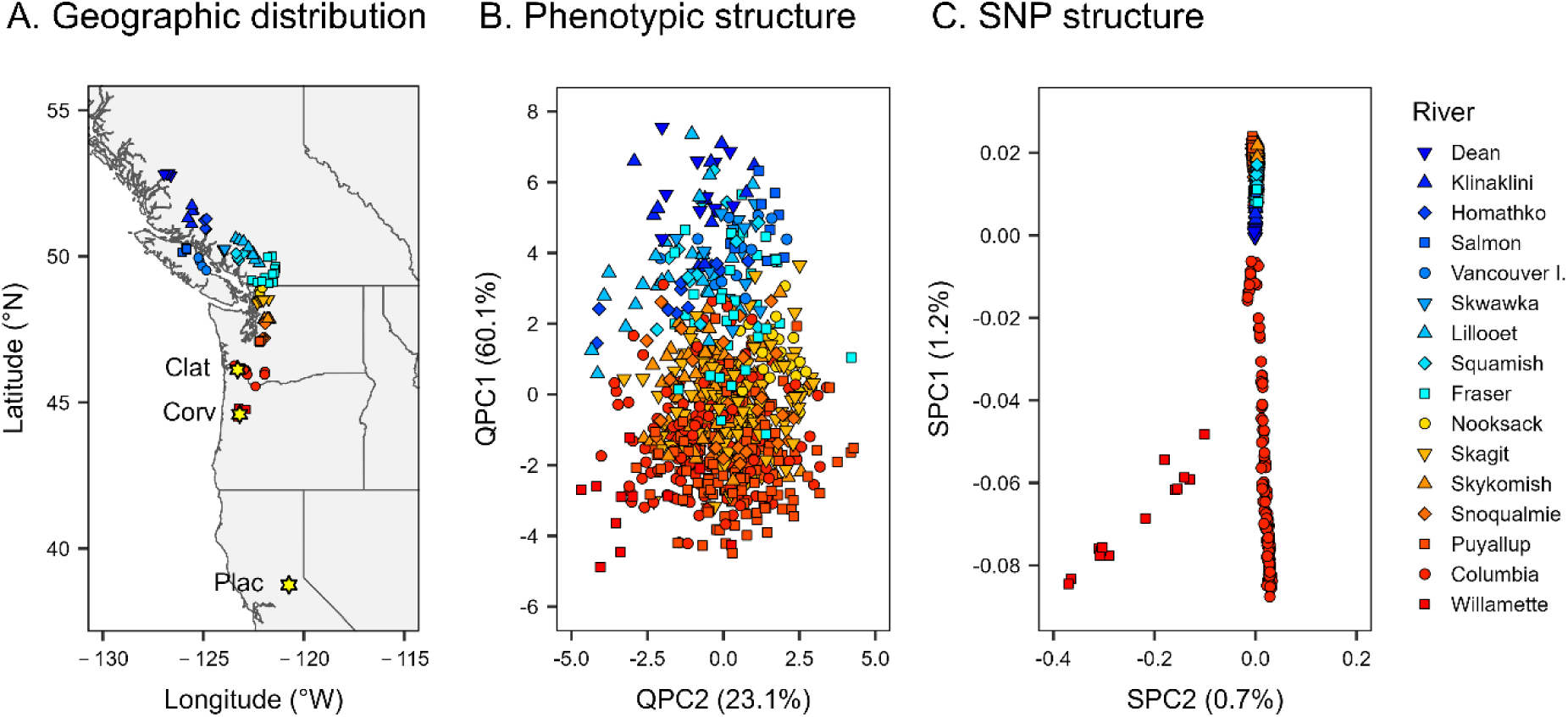
Geographic distribution (A), phenotypic population structure (B), and SNP population structure (C) for 840 *P. trichocarpa* clonal genotypes. (A) Source locations are color-coded by river with yellow stars indicating the locations of the three test plantations. (B) PC scores (QPC1 and QPC2) from the first two eigenvectors from a principal components analysis (PCA) of bud flush, bud set, and height phenotypes. (C) PC scores (SPC1 and SPC2) from the first two eigenvectors from a PCA of single-nucleotide polymorphism (SNP) markers filtered using ‘liberal’ criteria (*SI Appendix*, Table S4). The values in parentheses are the proportions of total variation accounted for by each PC score based on nine phenotypic variables (B) or SNP genotypes (C) from 840 clonal genotypes.

**Figure 2.**
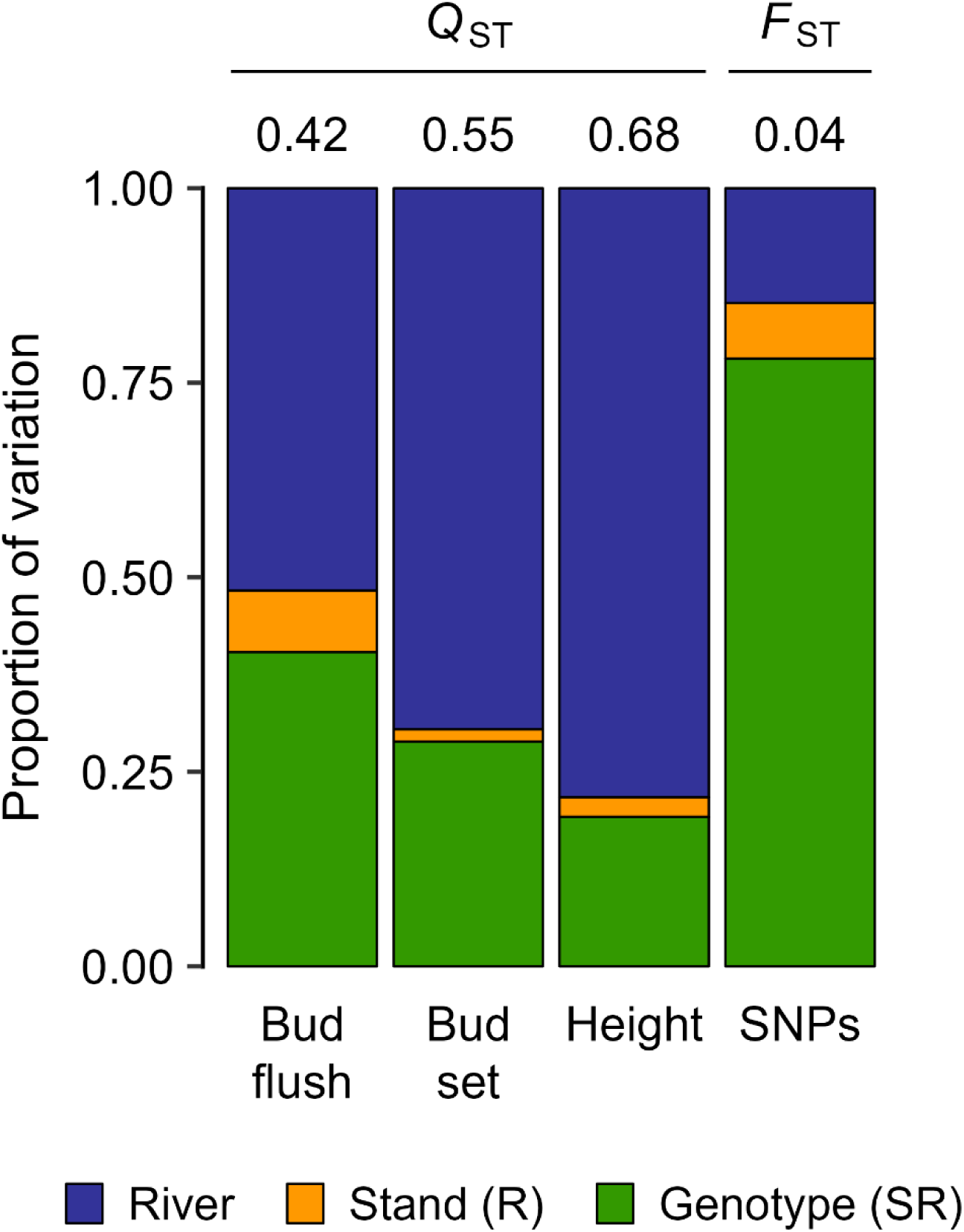
Distributions of genetic variation among rivers [River], stands-within-rivers [Stand (R)] and genotypes-within stands-and-rivers [Genotype (SR)] for 840 *P. trichocarpa* clonal genotypes. The y-axis shows relative proportions of variation based on mixed model analyses of quantitative traits and AMOVA for SNPs. Among-stand *Q*_ST_ values are shown above the bars for three quantitative traits (bud flush, bud set, and height), and the among-stand *F*_ST_ value is shown above the bar for SNPs.

### SNP variation was moderately structured and strongly associated with climate

Although SNP variation had a clear spatial pattern (Fig. 1C), the first two principal components explained only 1.9% of the total SNP variation among the 840 clonal genotypes, and the correlation between the first principal component for SNPs (SPC1) and latitude was moderate (r = 0.55). In contrast to *Q*_ST_, SNP differentiation was much lower (*F*_ST_ = 0.04, Fig. 2), but varied substantially among SNPs. Based on a sample of 1.1M SNPs, *F*_ST_ values were as high as 1.00 and the 99th percentile was 0.32. Thus, among all 20.8 M SNPs, there were at least 200K SNPs with very large *F*_ST_ values. Overall, variation among rivers accounted for 15% of the SNP variation (Fig. 2, fourth blue bar). Finally, variation for the first five SNP PC scores (SPC1-SPC5) was strongly associated with climate (*SI Appendix*, Tables S1 and S2).

### Adaptive trait phenotypes were predicted using geography, climate, or SNPs

We compared phenotypes predicted from geographic variables, climate variables, and SNPs using predictive ability (PA). PA is the Pearson product-moment correlation between the phenotypes described above versus phenotypes predicted from the measured trees using phenotypic BLUP (PBLUP). The overall ability to predict phenotypes across stands and rivers was moderate to high (PA > 0.5) using ridge regression with geography, climate, or SNP variables as predictors (Figs. 3 and 4A; *SI Appendix*, Table S3). Hereafter, we refer to ridge regression using SNPs or simulated RAD-Seq markers as genomic BLUP (GBLUP). On average, GBLUP PAs (0.702 to 0.735) were only modestly higher than PAs based on geography (0.659) or climate (0.683) (*SI Appendix*, Table S3). The absolute advantage of GBLUP was largest for bud flush. For this trait, GBLUP had a PA of 0.598 to 0.635. In contrast, the PAs were 0.531 based on geography and 0.579 based on climate variables.

**Figure 3.**
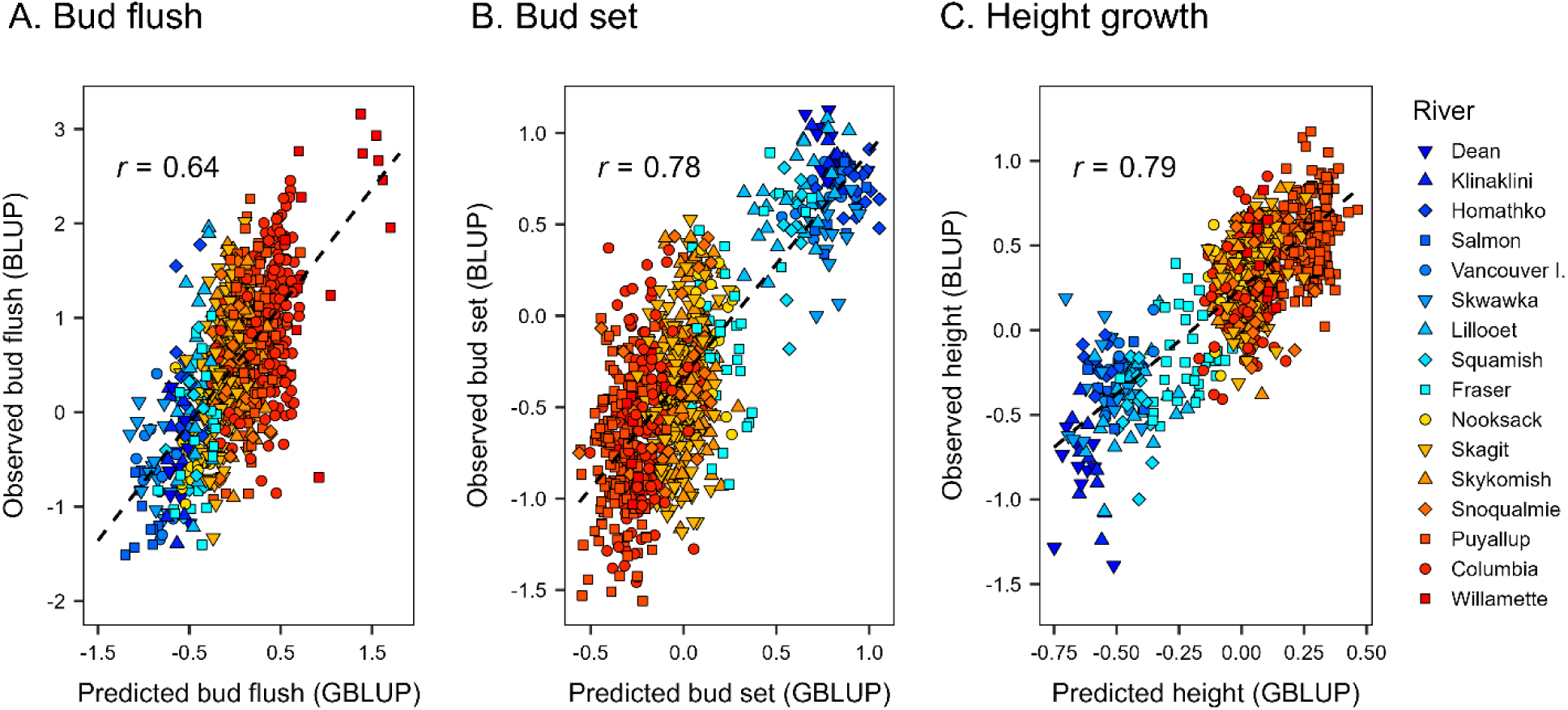
Predicted phenotypic values across all hierarchical levels (genotypes, stands, and rivers) based on field measurements (BLUP) versus SNPs (GBLUP). Predicted values for bud flush (A), bud set (B) and height growth (C) were based on field measurements of 840 *P. trichocarpa* clonal genotypes or single-nucleotide polymorphism (SNP) data. GBLUP values are averages of 100 random, 10-fold cross-validations with training population sizes of 756, prediction population sizes of 84, and 20,770,783 SNPs filtered using the ‘liberal’ criteria (*SI Appendix*, Table S4). The dashed line is the simple linear regression of BLUP on GBLUP.

**Figure 4.**
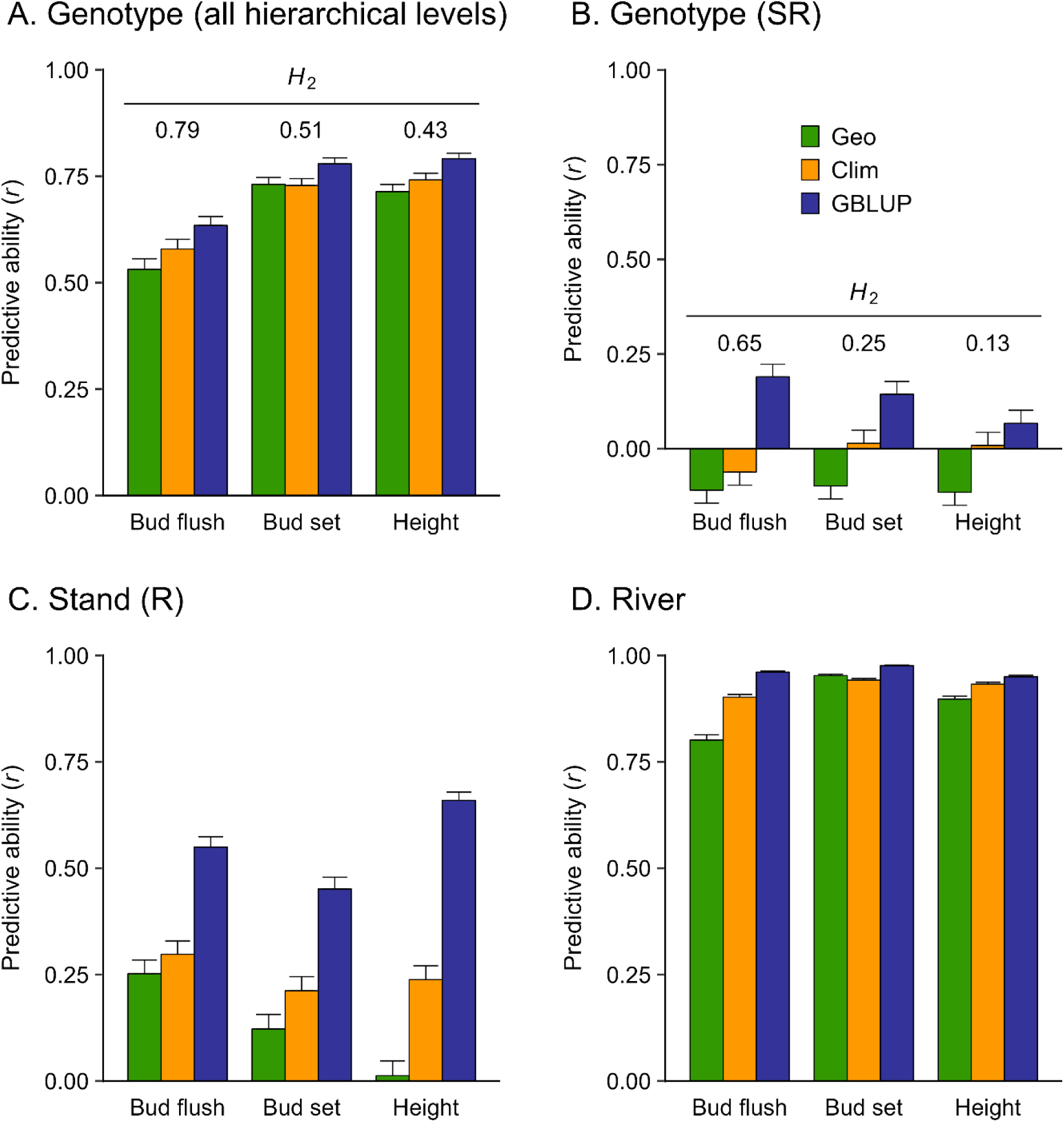
Predictive abilities (PA) for genotypes across all hierarchical levels (A), genotypes-within-stands-and-rivers (B), stands-within-rivers (C), and rivers (D). We used ridge regression with 3 geographic variables (Geo), 21 climatic variables (Clim), and 20,770,783 SNPs (GBLUP) filtered using the ‘liberal’ criteria (*SI Appendix*, Table S4). Bars are averages from 100 random, 10-fold cross-validations using training population sizes of 756 and prediction population sizes of 84. Standard errors (error bars) were calculated as described in the Materials and Methods. Broad-sense heritabilities (*H^2^*) are shown above the bars in A and B.

### Predictive ability was low after accounting for population structure

Next, we evaluated PA after rigorously accounting for population structure. Although PAs were high across the entire population, the clustering of genotypes by river (Figs. 1B-C, Fig. 3) suggested that much of the PA resulted from population structure. By partitioning the PAs into hierarchical levels (Figs. 4B-D), three important observations emerged. First, although PAs were moderate to high across the entire population (Fig. 4A), none of the models performed well within stands (PA < 0.2, Fig. 4B), even though 19-40% of the overall genetic variation occurred within stands (Fig. 2). Second, GBLUP models based on SNPs were consistently better at predicting stand-level phenotypes than were models based on geography or climate (Figs. 4A-D). Finally, the predictive abilities for river-level phenotypes were high for each model and trait (mean PA = 0.924, range = 0.801-0.976; Fig. 4D). These results are generally consistent with the hierarchical distribution of genetic variation for phenotypic traits, although within-stand PAs were disproportionately low (Fig. 4B) compared to the within-stand variance components (Fig. 2, first three green bars).

The low within-stand PA suggested that genomic prediction was affected by differences in genetic architecture among rivers. To test this, we developed GBLUP models using a subset of data from three well-sampled rivers. Specifically, we compared GBLUP models developed using genotypes sampled within rivers versus across rivers. The PAs of the within-river models (Skagit, Puyallup, and Columbia) were mostly larger or much larger than the PAs of the across-river models (Core and All), irrespective of training population size (Fig. 5). For bud flush, the PAs for the within-river models were more than twice as large as the PAs across rivers or across the entire study (Fig. 5A). We saw similar trends for bud set and height growth, but with some anomalies (Fig. 5B-C). Because training population size affects PA (Fig. 5, *SI Appendix*, Fig. S1), PAs may have been much higher and more consistent if we had more genotypes per river. For example, PAs increased with increasing training population size in the analyses described above (Fig. 5).

**Figure 5.**
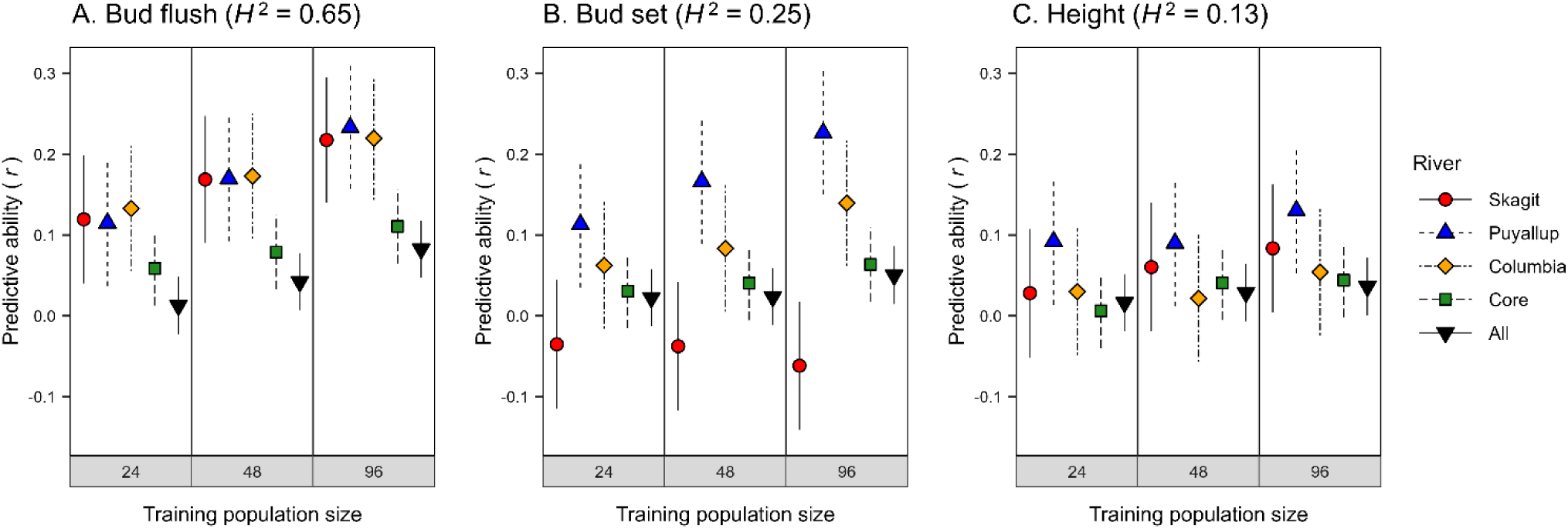
Predictive ability (PA) as a function of training population size using subsets of genotypes from the Skagit, Puyallup, and Columbia rivers. GBLUP analyses were conducted for genotypes-within-stands-and-rivers [Genotype (SR)] using SNP markers filtered using the ‘liberal’ criteria (*SI Appendix*, Table S4). PA was calculated using 160 clonal genotypes randomly selected from each of three rivers (Skagit, Puyallup, and Columbia), across all three rivers (Core), or from the entire population (All). Training population sizes were 24, 48 and 96, with a fixed prediction population size of 64 and all 20,770,783 SNPs. Averages were based on 100 replications of each analysis and standard errors (vertical lines) were calculated as described in the Materials and Methods. Genotype (SR) broad-sense heritabilities (*H^2^*) are shown in parentheses.

### Few SNP-phenotype associations were detected after accounting for population structure

To maximize the probability of detecting SNP-phenotype associations, we used 20.8M SNPs to conduct GWAS at two hierarchical levels; within-stands and across-stands-and-rivers.

These 20.8M SNPs were those remaining after using the ‘liberal’ filtering criteria (*SI Appendix*, Table S4). Using analyses designed to account for SNP population structure and cryptic relatedness (KANG et al. 2010), we detected many associations when we used phenotypes that incorporated variation among genotypes, stands, and rivers (Fig. 6A). However, when we conducted the same analyses within stands (i.e., using the genotype- within-stand phenotypes), we detected only one bud flush association (Fig. 6B). Results were similar when we used the first five PC scores (SPC1-SPC5) to correct for population structure, and when we excluded SNPs with minor allele frequency (MAF) < 0.01 (*SI Appendix,* Fig. S2A-B). In contrast, there was little difference between the two types of analyses (i.e., across all hierarchical levels versus among genotypes-within-stands) when we analyzed a less structured subset of genotypes from the Skagit, Puyallup, and Columbia Rivers (*SI Appendix,* Fig S2C). In both cases, only the single bud flush association was detected.

**Figure 6.**
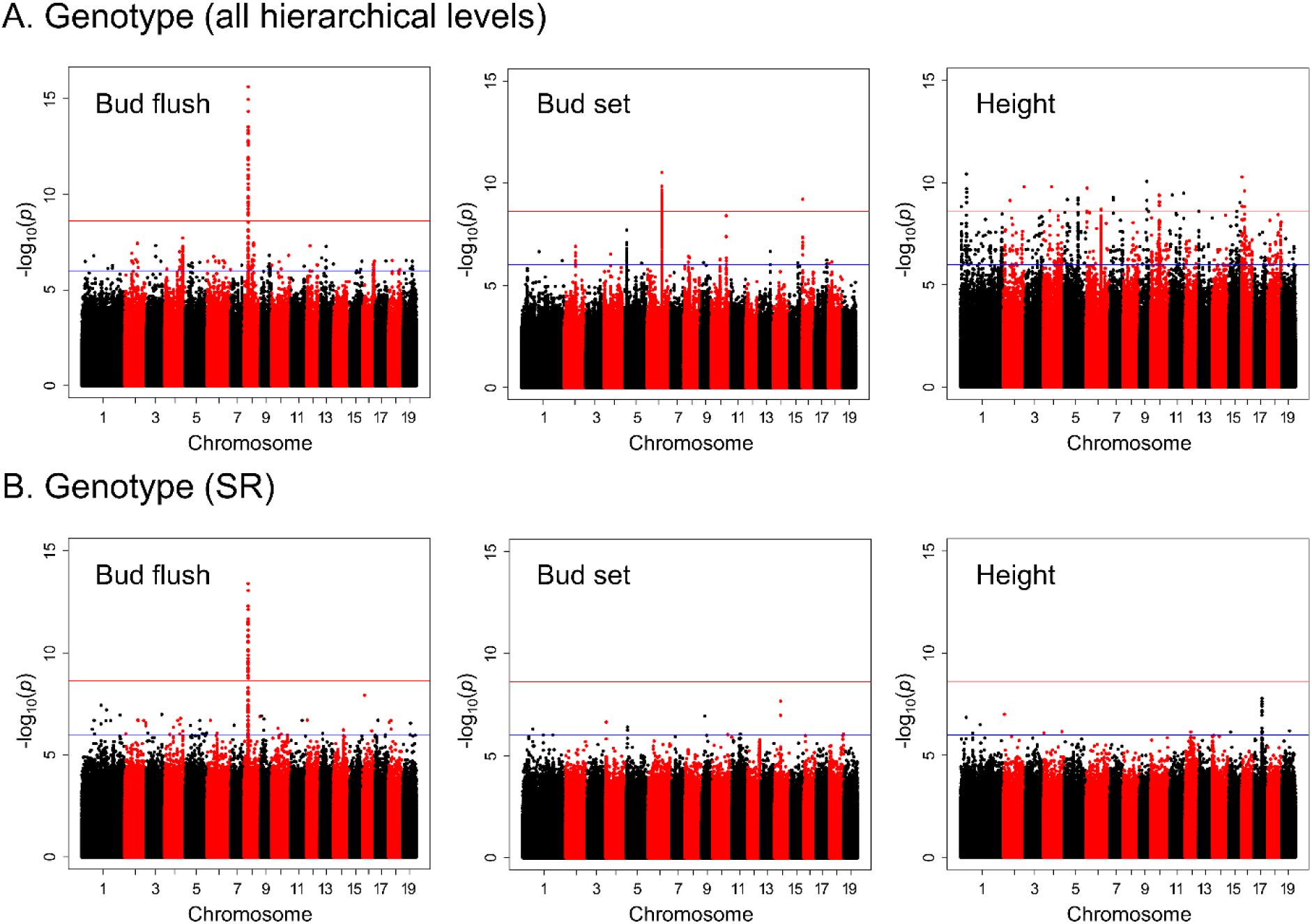
SNP-phenotype associations for bud flush, bud set, and height growth. A genome-wide association study (GWAS) was conducted across all hierarchical levels (A) and for genotypes-within-stands-and-rivers (B). GWAS was conducted using 20,770,783 SNPs filtered using the ‘liberal’ criteria (*SI Appendix*, Table S4) and the identity-by-state (IBS) kinship matrix. The blue line indicates a *P*-value of 10^-6^ and the red line indicates a Bonferroni-corrected *P*-value of 2.4 × 10^-9^ (α = 0.05).

### Within-stand genomic prediction and GWAS were probably limited by population differences in genetic architecture

We hypothesized that population differences in genetic architecture were partly responsible for the low PAs and few SNP associations within stands. To test this, we examined two important components of genetic architecture—allele frequencies at causal loci and linkage disequilibrium. Population differences in other components of genetic architecture, such as allele effect sizes and epistasis, may have also contributed (KACHURI *et al*. 2024) but were beyond the scope of this study because of the much larger sample sizes required for their reliable estimation.

First, we considered population differences in allele frequencies at causal loci (e.g., QTN or quantitative trait nucleotides). Allele frequencies affect the percentage of variation explained by a locus (PVE) and, thus, the ability to detect SNP-phenotypic associations using GWAS or predict phenotypes using GBLUP. Because we tested 20.8M SNPs (one SNP every 20 bp, *SI Appendix, Table S4*), our analyses likely included most of the QTN that explain the majority of variation for adaptive traits (i.e., excluding rare QTN with fewer than five minor allele copies, MAF < 0.003). As described above, allele frequency differences among rivers were substantial. Many pairwise *F*_ST_ values among rivers were close to 0.10 or greater (*SI Appendix,* Fig S3A), and allele frequency differences among rivers averaged 0.05 (SD = 0.06). Finally, differences in allele frequencies were structured—17% of SNPs (i.e., > 3M SNPs) had allele frequencies that were correlated with latitude at the river level (i.e., *P* < 0.05). Thus, not accounting for the substantial variability of causative locus PVE among rivers probably affected pooled within-stand GWAS and genomic prediction.

Second, we studied population differences in LD, which may result from differences in demographic history, allele frequencies, or linkage phases between loci. Consistent with previous results (SLAVOV *et al*. 2012), the extent of LD (i.e., average *r*^2^ > 0.2) varied roughly three-fold among rivers (6-18 kb) and tended to be higher within rivers than across rivers (*SI Appendix*, Fig. S4). The relationship between LD and physical distance varied substantially by MAF (Fig. 7A). Thus, we also quantified LD in MAF bins chosen to ensure all pairs of loci in a bin could have an *r*^2^ of at least 0.5 (*SI Appendix*, Table S5). In these analyses, LD was near zero for bins containing rare SNPs (MAF < 0.01). In contrast, for common SNPs (MAF ≥ 0.10), LD extended from 2-3 to over 10 kb (Fig. 7B). Because *r*^2^ values were highly variable, even within MAF bins (Fig. 7C), we estimated the probability that a causative polymorphism (e.g., QTN) would be tagged (*r*^2^ ≥ 0.6) by at least one SNP within 10 kb (Fig. 7D). This probability was sensitive to MAF filtering and the number of SNPs used to tag the QTN. Overall, more than 1M SNPs would be needed to tag a QTN with a probability of 0.5. Many more SNPs would be required to tag QTNs with even greater confidence, particularly if the QTN allele is rare (Fig. 7D). Thus, allele frequency differences among stands and rivers resulted in corresponding differences in LD, which likely affected the within-stand GBLUP and GWAS analyses.

**Figure 7.**
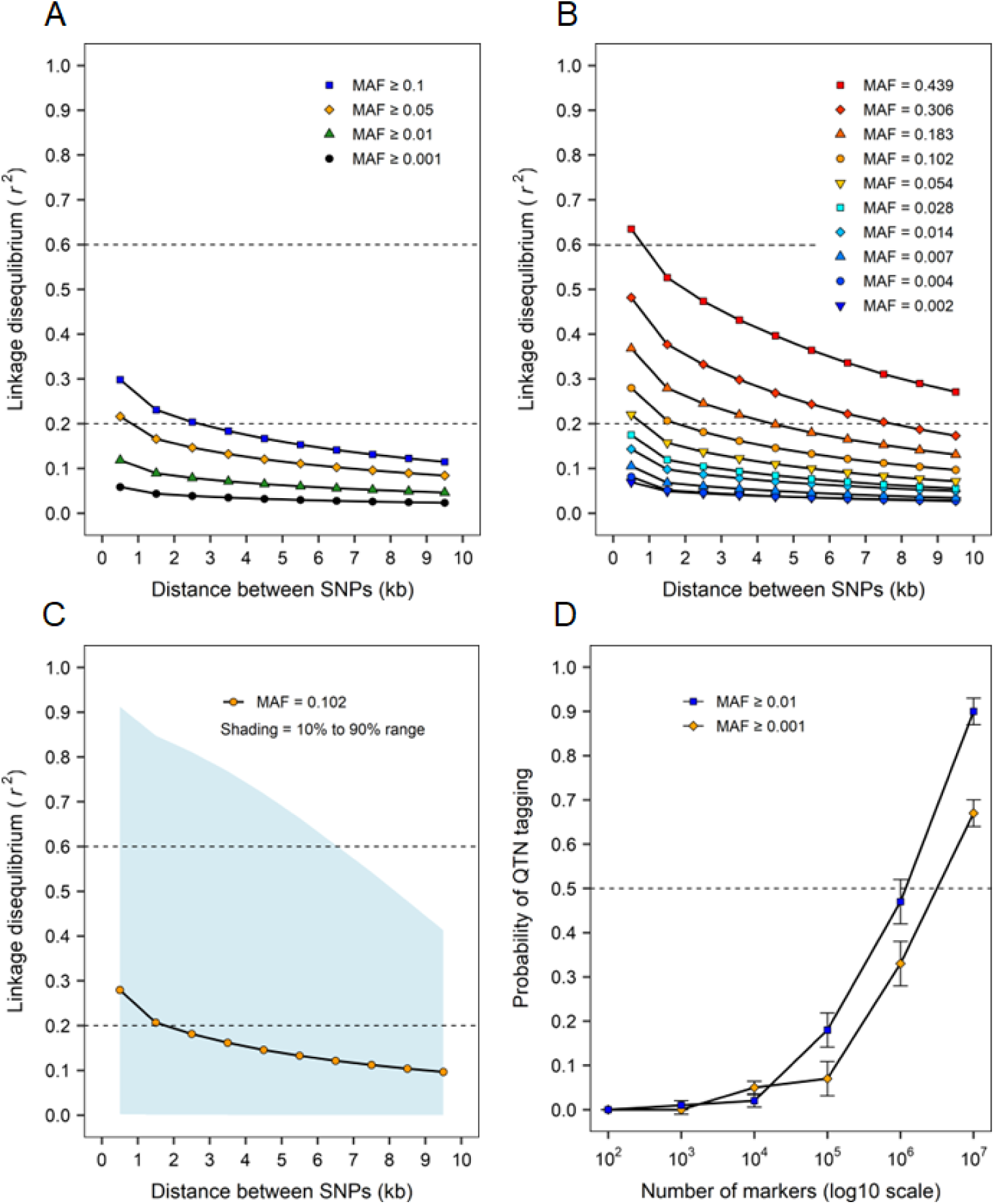
Linkage disequilibrium (LD) and probability of tagging causative loci using different minor allele frequency (MAF) thresholds. LD was calculated using different MAF filtering criteria (A), by MAF bin (B), and for a bin with MAF ranging between 0.071 and 0.132 (midpoint = 0.102) (C). MAF bin ranges (Table S4) are represented by their midpoints, and LD was calculated as the average *r*^2^ for pairs of SNPs in each 1-kb distance class. The probability of tagging a hypothetical quantitative trait nucleotide (QTN) was calculated using different numbers of randomly selected SNPs (D). Tagging was defined as the presence of at least one SNP in linkage disequilibrium (*r*^2^ ≥ 0.6) with the QTN within 10kb. Averages and standard errors (error bars) were based on 100 random samples.

Differences in linkage phase between QTN and linked markers, such as those resulting from different population histories, will also contribute to population variation in LD. To evaluate this contribution, we calculated haplotype sharing, a measure of linkage phase and allele frequency consistency among individuals, either within or among populations. Haplotype sharing was much lower for individuals sampled in different rivers than for individuals sampled in different stands-within-rivers and for individuals sampled within the same stand (Fig. 8A). Patterns of haplotype sharing for pairs of rivers strongly resembled those for allele frequency differentiation (*SI Appendix,* Fig. S3, *r* = -0.87, *P* < 10^-6^ from a Mantel test), suggesting that allele frequency differences were largely responsible for the reduced haplotype sharing among rivers. Indeed, haplotype sharing was much higher when analyses were limited to SNPs that had similar allele frequencies in each river, but the difference among hierarchical levels was still detectable (Fig. 8B).

**Figure 8.**
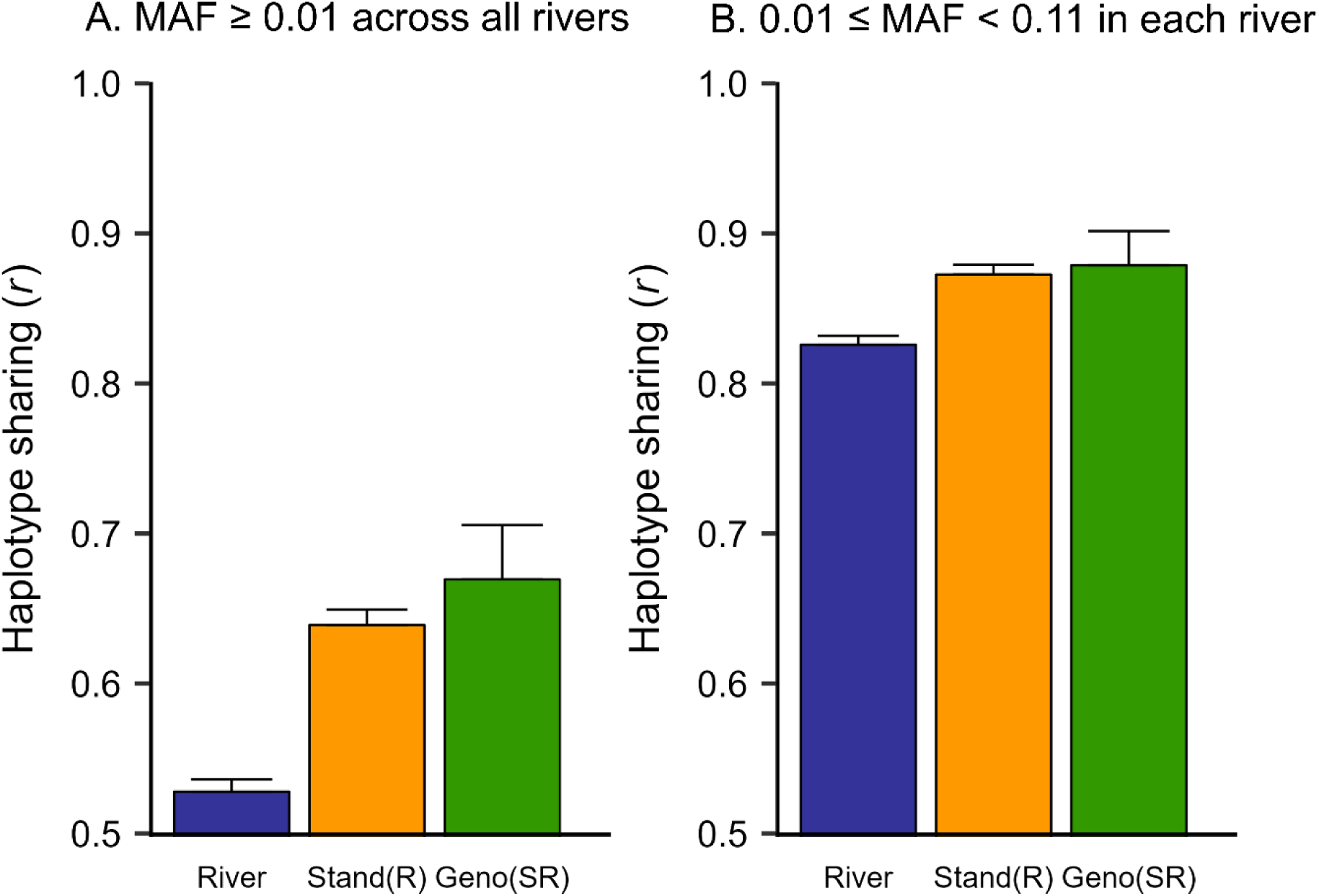
Linkage phase consistency (haplotype sharing) among genotypes-within- stands-and-rivers [Geno (SR)], stands-within-rivers [Stand (R)] and rivers [River]. (A) Analyses were based on 1,082,633 SNPs with MAF ≥ 0.01 separated by at least 300 bp and haplotype sharing was calculated for all pairs of SNPs located within 10 kb of each other. (B) Analyses were based on 1002 SNPs with 0.01 ≤ MAF < 0.11 in each of the 16 rivers and haplotype sharing was calculated for all pairs of SNPs located within 1 Mb of each other. Standard errors (error bars) were calculated as described in the *SI Appendix,* Materials and Methods.

### Delineation of seed zones

To illustrate the practical implications of our results, we compared the ability of different types of data (i.e., phenotypic, geographic, climate, and SNPs) to reconstruct seed deployment zones delineated using different criteria (Fig. 9). Seed zones are geographic areas of genetic and environmental homogeneity used to guide deployment (e.g., planting) of tree genotypes (HOWE *et al*. 2006). For natural populations, we assume that genotypes collected within a seed zone can be deployed within the same zone without risking maladaptation. When the ‘true’ seed zones were assumed to correspond to the 16 rivers (Fig. 9A), phenotypic data were best for reconstructing these zones, although SNPs were only slightly less accurate (cluster purity of 0.758 vs 0.714). Purity is a 0 to 1 measure of cluster quality, or the extent to which a clustering method recovers known classes. The cluster purities of geographic and climate data were substantially lower for this scenario (0.626 and 0.571). However, when true seed zones were delineated based on phenotypic data, reconstructions based on geographic, climate, and SNP data had similar cluster purities (Fig. 9B, C).

**Figure 9.**
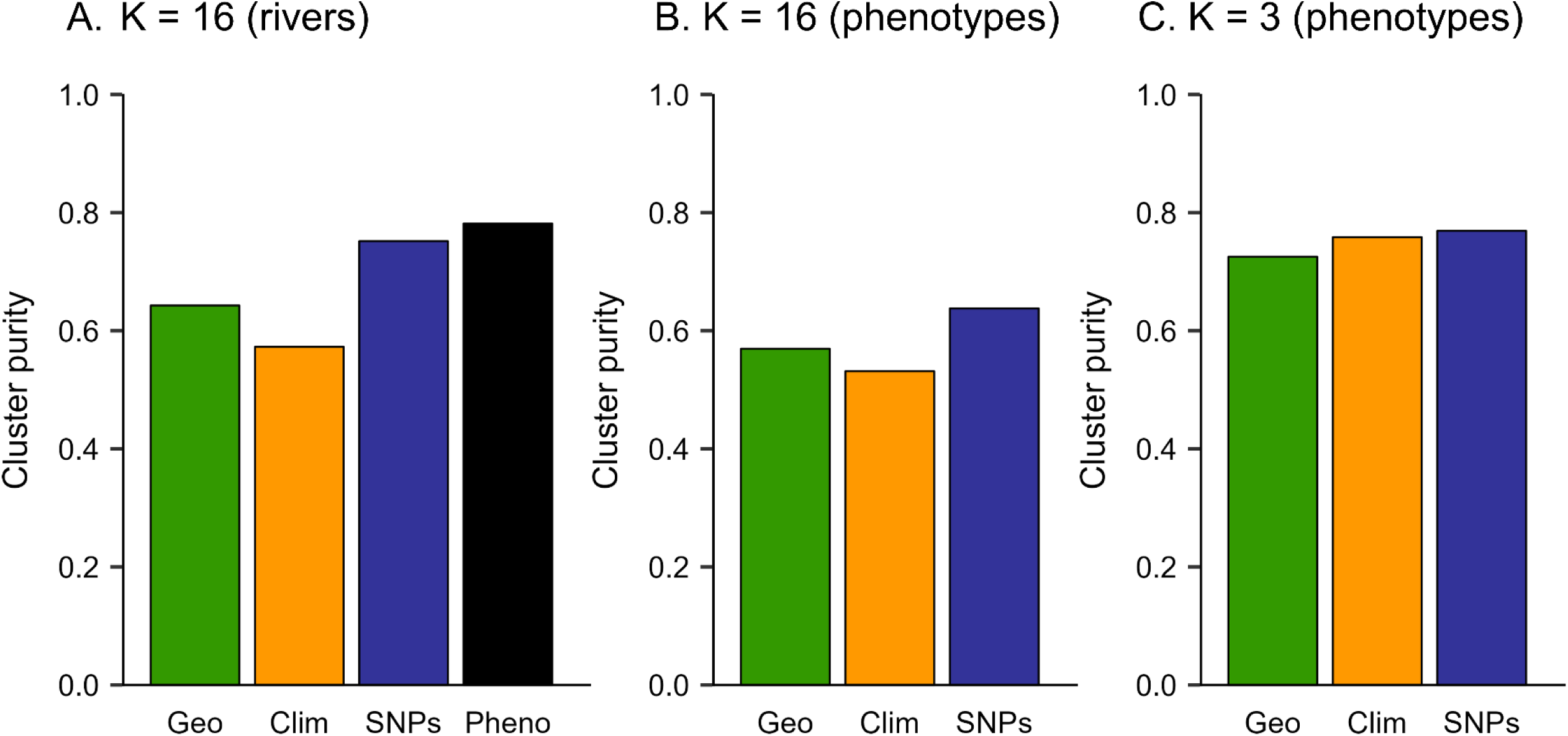
Delineation of stand-level seed zones using geographic (Geo), climate (Clim), SNP (SNPs), or phenotypic data (Pheno) for 840 *P. ichocarpa* clonal genotypes sampled from 91 stands in 16 rivers. Different numbers (K) of ‘true’ seed zones were assumed to correspond to rivers A), or seed zones were defined using k-means clustering of phenotypes (B, C). The success of reconstructing true seed zones based on different types f data was evaluated using cluster purity, which is the proportion of stands that were both in the same true seed zone and in the same reconstructed seed zone. Averages are based on 100 replications of each analysis.

## Discussion

Our results highlight the challenges of using genomic information to understand the genetics of complex quantitative traits in natural populations. Because climate is a primary driver of local adaptation (WADGYMAR *et al*. 2022), we focused on climate adaptation traits, or simply ‘adaptive traits.’ These traits have been important in tree breeding programs (HOWE *et al*. 2006; MACLACHLAN *et al*. 2017; HUREL *et al*. 2021) and for guiding assisted migration to respond to climate change (PARK AND TALBOT 2018). We studied height growth and vegetative bud phenology using GWAS and genomic prediction, two widely used approaches in forest trees (GRATTAPAGLIA *et al*. 2018; GRATTAPAGLIA 2022; STRAUSS *et al*. 2022).

GWAS has been used in natural populations—with the goal of identifying causal loci (EVANS *et al*. 2014; DE LA TORRE *et al*. 2021), enhancing tree breeding (HIRAOKA *et al*. 2018; MÜLLER *et al*. 2019) or informing assisted migration (RELLSTAB *et al*. 2021). Nonetheless, many studies are compromised by population structure, the inclusion of closely related individuals, low marker coverage, or small population sizes (i.e., low power) (WEISS *et al*. 2020; STRAUSS *et al*. 2022). These limitations are likely to lead to the misidentification of causal loci and other misinterpretations. In contrast, genomic prediction works best in populations of closely related trees and the identification of causal loci is not an explicit goal. Thus, genomic prediction has been used to select desirable genotypes in breeding populations of forest trees (reviewed in GRATTAPAGLIA 2022). We partitioned genetic variation into hierarchical levels (i.e., river, stand-within-river, and genotype-within-stand) to evaluate GWAS and genomic prediction for understanding complex quantitative traits in natural populations of black cottonwood. These analyses highlight the challenges of using GWAS and genomic prediction across populations with different genetic architectures. Finally, compared to genomic information, population-level phenotypes were predicted nearly as well by climate alone.

### Genomic prediction and GWAS were highly sensitive to population genetic structure

We showed that phenotypic variation in adaptive traits was highly structured and strongly associated with climate, which is consistent with other studies of black cottonwood and other wide-ranging tree species (PAULEY AND PERRY 1954; WEBER *et al*. 1985; DUNLAP AND STETTLER 1996; GORNALL AND GUY 2007; ST CLAIR AND HOWE 2007; ALBERTO *et al*. 2013; EVANS *et al*. 2014; MCKOWN *et al*. 2014a; PORTH *et al*. 2015; HOLLIDAY *et al*. 2016; FRANK *et al*. 2017; MCKOWN *et al*. 2018). In contrast, SNP variation was moderately structured but clearly associated with climate. Other population genomic studies found similar evidence for SNP population structure, but patterns of variation were typically much weaker than for adaptive traits, both in *Populus* (SLAVOV AND ZHELEV 2010; EVANS *et al*. 2014; MCKOWN *et al*. 2014a; PORTH *et al*. 2015) and other trees (HOWE *et al*. 2003; ALBERTO *et al*. 2013). In our study, the difference between phenotypes (average *Q*_ST_ = 0.55) and SNPs (overall *F*_ST_ = 0.04) was pronounced but not surprising, given that most SNPs are expected to be selectively neutral.

When we conducted analyses across all hierarchical levels (i.e., rivers, stands, and genotypes), we detected one association with bud flush, two associations with bud set, and over a dozen associations with height growth. However, when we accounted for population structure using PCs (PRICE *et al*. 2010; EVANS *et al*. 2014) and by conducting within-population analyses, most associations disappeared. This suggests they were either false positives caused by population stratification or true associations that were strongly correlated with population structure, and their further prioritization would require strong support from other lines of evidence (FURCHES *et al*. 2019; CHHETRI *et al*. 2020). However, a single bud flush association remained significant, even after excluding rare alleles (i.e., MAF < 0.01) and analyzing a sub-set of genotypes from three rivers in the core of the species range. This bud flush association, which included 30 common SNPs (MAF ∼ 0.10, p-value < 2.4 × 10^-9^) in strong LD (see below) spanning a region of nearly 60 kb, was also reported by EVANS *et al*. (2014). In contrast, MCKOWN *et al*. (2018) found no bud flush associations in this region using GWAS on black cottonwoods sampled mostly from British Columbia. This difference highlights the influence of genetic architecture on the ability to detect causal loci using GWAS (discussed below). In GWAS, uncorrected population structure will likely lead to misidentification of causal loci, and the numbers of false positives can become very large as more SNPs are analyzed. For example, among the 20.8M SNPs we analyzed, ∼3.5M had allele frequencies that were significantly correlated with latitude (uncorrected p-value = 0.05), which may lead to situations where even rigorous accounting for population structure may fail (PRICE *et al*. 2010). Thus, because adaptive traits tend to correlate with population structure, only associations that are also detected within populations can be considered robust.

Likewise, our ability to predict phenotypes using SNP markers mostly resulted from population structure—PAs were high at the river level (0.950-0.976), moderate at the stand- within-river level (0.451-0.659), and low for genotypes within stands (0.067-0.190). Instead of population structure, within-stand prediction presumably resulted from linkage between SNPs and causal loci or the ability of SNPs to estimate relatedness among trees. For breeding programs, we see a limited role for genomic prediction in natural populations, at least for adaptive traits. First, although genomic prediction worked well across populations, prediction was almost as good using climate variables alone (see below). Second, predictive abilities were low within populations. Hypothetically, genomic prediction in natural populations could be practically useful in breeding if (1) most of the selection intensity is allocated to a spatial scale where population structure is not sufficiently well captured by climate data and (2) a substantial proportion of phenotypic variation occurs at that spatial scale. For example, genomic predictions at the stand-within-river level were much better than predictions from climate (Fig. 4C). However, only 2-8% of the genetic variation for adaptive traits occurred at the stand-within-river level (Fig. 2).

### Population-level phenotypes can be predicted using SNPs or climate variables

There has been much interest in using genomic information to infer maladaptation to future climates and guide assisted migration (BORRELL *et al*. 2020; MACLACHLAN *et al*. 2021; RELLSTAB *et al*. 2021). Thus, a variety of statistical approaches have been developed to predict maladaptation from genomic data (e.g., genomic offsets, RELLSTAB *et al*. 2021) that are analogous to earlier approaches using phenotypes (CAMPBELL 1986; ST CLAIR AND HOWE 2007; FRANK *et al*. 2017). Using phenotypes measured in the field, we demonstrated that climate adaptation traits can be predicted using SNPs but were predicted nearly as well using climate variables alone. Furthermore, there were few differences between seed zones delineated using phenotypes, SNPs, or climate variables. Genomic offset experiments suggest that SNP-based genomic offsets can be used to predict population phenotypes better than climate or geographic variables alone, although not consistently (FITZPATRICK *et al*. 2021; LIND *et al*. 2024). Simulations suggest that *a priori* selection of climate variables improves climate-only models (LIND AND LOTTERHOS 2024), and empirical studies of genomic offsets indicate that predictions can be improved further by using SNPs to select/weight climate variables or infer non-linear relationships between climate distance and phenotypes (FITZPATRICK *et al*. 2021; LÁRUSON *et al*. 2022). However, the most important (i.e., selective) climate variables were not always identifiable in simulations (LÁRUSON *et al*. 2022; LIND AND LOTTERHOS 2024).

As we found using genomic prediction, the performance of genomic offsets seems to rely on population structure—random markers performed as well as known causal markers in simulations (LIND AND LOTTERHOS 2024) and as well as candidate loci in empirical studies (FITZPATRICK *et al*. 2021; LIND *et al*. 2024; but see GAIN *et al*. 2023). In addition to being surrogates for phenotypic population structure, SNPs may enhance prediction by reducing error in climate variables. Predictions from climate interpolation models are not without error, particularly in remote and mountainous regions and when low resolution climate data are used (e.g., 1 x 1 km, FITZPATRICK *et al*. 2021; LIND *et al*. 2024)—and adding SNP data may counteract these errors. If so, climate-only models might be improved by increasing the accuracy of climate data; perhaps by establishing networks of ‘micro’ weather stations (YE *et al*. 2022). Because this would improve assisted migration for all species, it might be a wiser use of resources compared to developing new genomic resources for many individual species. Second, SNPs might improve predictions by accounting for phenotypic relationships among populations unrelated to climate—e.g., those resulting from demographic processes such as colonization, migration, or secondary contact. In any case, based on our results and genomic offset studies, it is unlikely that the predictive power of genomic offsets comes from information derived from causal loci. On the other hand, using SNPs to guide assisted migration has two potential pitfalls. First, neutral population structure may follow different spatial patterns compared to phenotype-climate associations, leading to poor prediction of maladaptation. Second, because the acquisition of SNP data will probably delay assisted migration for most species, it might be more pragmatic to use climate-only models instead.

Thus, because phenotype-climate associations are reasonably well understood across species (HOWE *et al*. 2003; AITKEN AND BEMMELS 2016; LEITES AND GARZON 2023), we argue that important climate variables can be reasonably selected *a priori* and used to guide assisted migration in the absence of SNP data.

### Why were PAs so low and why did we find few GWAS SNPs within populations?

Given the large number of SNPs we used, why was it difficult to detect associations and predict phenotypes within populations (i.e., after rigorously accounting for population structure)? Based on our results and interpretation of the relevant literature, we offer four main explanations: (1) complex quantitative traits are controlled by many genes with small effects, (2) frequencies of causative polymorphisms differ among populations, (3) linkage disequilibrium is mostly low, particularly for rare alleles, and (4) frequencies of marker alleles and LD differ among populations.

A long history of pedigree-based QTL studies and GWAS analyses indicate most traits in forest trees are controlled by many loci with small effects (GRATTAPAGLIA *et al*. 2018). Furthermore, because of small sample sizes and clustered loci, effect sizes are probably overestimated, and the true number of loci is probably underestimated (BEAVIS 1998; HALL *et al*. 2016). Studies of outcrossing plants, livestock, and humans lead to similar conclusions (VISSCHER *et al*. 2012; BERNARDO 2016; VISSCHER *et al*. 2017). These factors have three important effects for complex quantitative traits. First, very large sample sizes will be needed to detect most small-effect loci, probably many more than have been used or perhaps are even feasible (VISSCHER *et al*. 2017). Second, low-powered experiments are likely to report many spurious associations. For example, the large number of associations reported in some recent studies of forest trees were considered ‘surprising’ based on theoretical expectations (STRAUSS *et al*. 2022), and others highlighted the lack of GWAS reproducibility (GRATTAPAGLIA *et al*. 2018). Finaly, thousands of GWAS loci may be needed to explain most of the genetic variation in quantitative traits. The challenges in understanding the genetic basis of human height provide a cautionary tale. Recent success at explaining most of the variation in human height (e.g., > 50%) required millions of study participants and more than 12 K independent GWAS loci (i.e., SNP associations) (YENGO *et al*. 2022).

Causal loci are difficult to detect because the frequencies of causative polymorphisms differ among populations (KACHURI *et al*. 2024). Because we used 20.8 M markers with an average spacing of about 20 nt across the genome, we probably genotyped most of the causal QTN that cumulatively explain most of the phenotypic variance, yet detected few GWAS loci. The ability or power to detect a causal locus (*c*) depends on sample size (*N*) and the proportion of phenotypic variance explained by the causal locus (*PVE*_*c*_ = *V*_*g*_⁄*V*_*t*_), where *V*_*g*_ is the genetic variation explained by the locus and *V*_*t*_ is total genetic variation. Furthermore, *PVE*_*c*_ is a function of allele effects and MAF. For example, when allele effects are estimated by the standardized regression coefficient (β) of phenotype on allele number (i.e., 0, 1, 2), *PVE*_*c*_ = 2 · β^2^ · *MAF*(1 − *MAF*) (Visscher et al 2017). Because allele effects are unlikely to vary much among closely related populations (WANG *et al*. 2020; KACHURI *et al*. 2024), the most important contributors to GWAS power are *MAF*, number of contributing loci (reflected in *V*_*t*_ and the standardized regression coefficient), and experimental *N*. Thus, when MAF varies across populations, GWAS power also varies, which contributes to poor reproducibility. Although pooled within-population analyses can account for the main effects of population structure, differences in MAF are still problematic, particularly for loci that are not segregating in some populations. Additionally, it is probably impossible to assay all causal loci using SNPs alone because phenotypic variation also arises from other types of genomic variation (LIN *et al*. 2024). Finally, population differences in LD (i.e., *r*^2^) reduce power even further when linked markers (*m*) are used to detect causal loci. In this case, *PVE*_*m*_ = *r*^2^ · *PVE*_*c*_ (Visscher *et al*. 2017).

Generally low LD in forest trees has historically been attributed to an outcrossing mating system, large effective population size, weak selection, and little population structure for most loci (NEALE AND SAVOLAINEN 2004; NEALE AND KREMER 2011). However, more recent studies revealed some notable exceptions (SHALEV *et al*. 2022), and generally indicate that LD is higher than previously thought and highly variable across the genome (SLAVOV *et al*. 2012; SILVA-JUNIOR AND GRATTAPAGLIA 2015; BUTLER *et al*. 2022). In our study, LD for common SNPs (i.e., MAF > 0.1) decayed below 0.2 within 1 to 3 kb on average, and extended well beyond 10 kb for more than 10% of SNP pairs (Fig. 7C). On the other hand, LD was near zero for rare SNPs (MAF < 0.01, Fig. 7A,B). Mostly low LD and small locus effect sizes make it difficult to identify causal loci using linked markers. Furthermore, GWAS power is particularly low when allele frequencies differ between markers and causal loci (WRAY 2005). In fact, WRAY (2005) concluded that constraints on allele frequencies are ‘severe’—for a causal locus with MAF of 0.1, the MAF of the marker must be within 0.02 for *r*^2^ to be ≥ 0.8. That is, strong constraints on allele frequencies make causal loci ‘invisible’ to most nearby markers. In our study, an LD > 0.6 seemed necessary to detect the single bud flush locus. Using 0.6 as the LD cut-off, more than 1M SNPs with MAFs > 0.01 would be needed to have a 50% chance of tagging a causal locus (Fig 7D). Overall, our ability to detect the bud flush locus seemed to rely on a high MAF for the causal locus (i.e., > 0.05), high heritability for the phenotypic trait (0.79), and large locus effect size (i.e., PVE ∼5%).

Population differences in linkage phase may also obscure species-wide associations using linked markers—SNPs associated with positive phenotypes in one population may be associated with negative phenotypes in another. This may have been a contributing factor in our study because haplotype sharing was greatest within stands and lowest among rivers (Fig. 8A). However, we showed that the reduced haplotype sharing among rivers was mostly due to differences in allele frequencies (Fig. 8B). Thus, differences in linkage phase are probably not the main reason for low GWAS power and predictive ability.

Overall, while other differences in genetic architecture (e.g., allele substitution effects or epistasis) may also be contributing, we hypothesize that the main limiting factors for GWAS and genomic prediction are allele frequency differences in causal and marker loci among populations (WANG *et al*. 2020; KACHURI *et al*. 2024). Across population analyses lead to incorrect inferences about the causal relationships between SNPs and phenotypes, whereas pooled within-population analyses have low power to detect GWAS loci or predict phenotypes.

### Implications

We show that across-population GWAS and genomic prediction are strongly influenced by population structure, rather than the causal relationships between SNP loci and adaptive traits. Thus, across-population analyses promote incorrect inferences about causal loci.

Instead, analyses of single-populations or the use of pooled within-population analyses should lead to more robust conclusions. The drawback of using a single population is that causal loci may be missed because they are not segregating. The drawback of using pooled within-population analyses is that power is compromised by differences in genetic architecture among populations. In any case, to detect most SNP-trait associations and predict phenotypes accurately, population sample sizes in the tens to hundreds of thousands will probably be needed. Obviously, experiments of this size will be infeasible for most forest tree species, even for a single population. Furthermore, based on human studies, substantially larger experiments may be needed (YENGO *et al*. 2022). Thus, we conclude that GWAS analyses are unlikely to detect most of the causal loci, explain a substantial proportion of trait heritability, or contribute meaningfully to traditional tree breeding, gene conservation, or assisted migration. GWAS can almost certainly be used to detect some of the causal loci, but ultimately transgenic or gene editing approaches may be required to distinguish candidate from causal loci (STRAUSS *et al*. 2022). Likewise, the success of within-population genomic prediction will improve as sample sizes become larger, but predictive abilities in most natural populations will always be constrained by the low relatedness among trees. Compared to the predictive abilities of progeny tests alone, it is questionable if genotyping and other costs needed to use genomic prediction in natural populations will be justified for adaptive trait breeding or assisted migration.

Despite the challenges, substantial research has been devoted to identifying or tagging causal loci for practical applications. In contrast, our results demonstrate the power of using neutral loci or climate variables to predict population-level phenotypes, at least for species with local adaptation to climate. These conclusions are supported by results from genomic offset simulations and validation experiments (discussed above). We hypothesized that SNPs improve climate-based prediction of population phenotypes by helping to characterize population structure, particularly when inappropriate climate variables are used or when the climate variables have error. Nonetheless, our results suggest that wise use of climate variables alone can be used to predict population phenotypes, delineate seed zones or deployment zones, and guide assisted migration.

## Materials and Methods

### Plant materials and test plantations

In the winter of 2008, we obtained or collected stem cuttings from 1101 *P. trichocarpa* genotypes, representing a large portion of the latitudinal range of the species (Fig. 1A; (SLAVOV *et al*. 2012). In April and May of 2009, we established rooted cuttings in three test plantations spanning the south-central portion of the black cottonwood range west of the Cascade Mountains (Fig. 1A, *SI Appendix,* Materials and Methods).

### Phenotypic measurements and analysis

Between 2009 and 2013, we measured height growth and two phenological traits, vegetative bud flush (BF) and bud set (BS), by visually classifying the phenological state of each tree using six-stage scoring scales (*SI Appendix*, Fig. S6). For each plantation and year, we chose measurement dates to maximize the phenotypic variation in BF and BS. In addition, we measured the current and previous year heights of the main stem as the distance from the groundline to the apical bud or to the most recent budscale scars (i.e., position of last year’s apical bud). For data analyses of height growth (HT), we averaged height growth for the 2010, 2011, and 2012 growing seasons. Similarly, when multiple BF and BS measurements were available, we first identified the measurement with the highest heritability for a given year (*SI Appendix,* Materials and Methods) and then averaged measurements across years.

Finally, we used mixed linear models to estimate variance components, heritabilities, genetic correlations, and random effects (i.e., best linear unbiased predictors, BLUPs) at the river (R), stand-within-river [S(R)], and genotype-within-stand-and-river [G(SR)] levels (*SI Appendix,* Materials and Methods). Thus, this approach allowed us to partition genetic variance and calculate “phenotypes” at three hierarchical levels (i.e., river, stand, and genotype), as well as across all levels (G).

### SNP data

We obtained data for 28,342,758 biallelic SNPs (https://cbi.ornl.gov/gwas-dataset/) from 970 *P. trichocarpa* individuals (clonal genotypes), and then removed 130 individuals from this dataset for the final analyses. We excluded individuals with a mean sequencing depth < 7, eliminated close relatives using an approach similar to that of EVANS *et al*. (2014), and then excluded 42 other individuals for other reasons (*SI Appendix*, Materials and Methods). The remaining 840 clonal genotypes represented 91 stands in 16 rivers (Fig. 1). For the final analyses, we used VCFtools v. 0.1.14 (DANECEK *et al*. 2011) and PLINK v.1.90b4.4 (CHANG *et al*. 2015) to filter SNPs based on ‘strict’ and ‘liberal’ criteria, and then simulated a set of 51,820 ‘RAD-Seq’ markers (*SI Appendix*, Materials and Methods and Table S4).

### SNP population structure and allele frequency differences

We calculated individual-tree PC scores using the liberally filtered SNP set and the SMARTPCA software package (v. 13050; PATTERSON *et al*. 2006). For this analysis, we selected a subset of non-singleton SNPs separated by at least 300 bp (*vcftools --thin* 300), and then removed one SNP from each pair of loci linked at *r^2^* ≥ 0.8 to avoid artefacts caused by large blocks of tightly linked markers (PATTERSON *et al*. 2006; NELSON *et al*. 2008).

Bivariate plots of PCs were used to reveal population structure at the river level. We also used SMARTPCA to calculate pairwise estimates of *F*_ST_ at the river level based on Hudson’s estimator, which is robust to the effects of rare-allele SNPs (BHATIA *et al*. 2013). Finally, we used the same SNPs and the HIERFSTAT package in R (GOUDET 2005) to estimate SNP variance components and hierarchical *F*-statistics at the river, stand, and genotype levels.

This analysis was designed to match our analyses of phenotypic data (see above, *SI Appendix,* Materials and Methods).

To quantify allele frequency differences among rivers, we first calculated the allele frequencies of all liberally filtered SNPs (*plink --freq --family*) in each river. Then, we calculated pairwise allele frequency differences among rivers and correlations between the river-level allele frequencies and latitude using R (R CORE TEAM 2021).

### GBLUP and GWAS

We used the *kin.blup* function of the rrBLUP R package (i.e., GBLUP approach; ENDELMAN 2011) to predict phenotypes based on SNP markers (i.e., GBLUP approach; MEUWISSEN *et al*. 2001; DE LOS CAMPOS *et al*. 2013). Genomic relationship matrices (GRMs) were calculated using the *kin.blup* function of rrBLUP or the --*make-grm-alg 1* option of GCTA (YANG *et al*. 2011). We also used a subset of analyses to test various Bayesian approaches implemented in the BGLR R package (PEREZ AND DE LOS CAMPOS 2014), and the results were essentially the same. The phenotypes for BF, BS, and HT were the random effects for three hierarchical levels, *G*(*SR*), *S*(*R*), and *R*, as well as the combined effects across all levels (*G*), using random effects from Model 2 (*SI Appendix*, Materials and Methods). We tested the effect of training population sizes (*N_t_*) ranging from 48 to 756, while keeping the prediction population size (*N_p_*) fixed at 84. This was done by randomly sampling *N_t_* + *N_p_* clonal genotypes, and then randomly allocating them to the training versus prediction population. Similarly, for river-specific GBLUP models, we varied *N_t_* between 24 and 96, while *N_p_* was fixed at 64. For all other GBLUP analyses, we used random, ten-fold cross- validation (*N_t_* = 756, *N_p_* = 84). To assess the effect of the number of markers used (*M*), we randomly selected 10^1^ – 10^7^ SNPs at random using PLINK (*plink --thin-count*). The performance of GBLUP was measured by predictive ability (PA), defined as the Pearson product-moment correlation coefficients between the input phenotypes and the phenotypes predicted from the SNP data (DAETWYLER *et al*. 2013; SLAVOV *et al*. 2014). For testing the effect of *N_t_*, we calculated PAs based on the 64 or 84 predicted values (*N_p_*), but for the analyses using 10-fold cross-validation, PAs were based on all 840 predicted values. We calculated PAs at the river and stand-within-river levels by calculating the correlation between the R and S(R) phenotypes versus the average predicted values from GBLUP for the clonal genotypes in each river or stand. Standard errors of PA means were calculated as [*var*(*r*)/*k* + (1 − *r*^2^)^2^⁄(*N* − 2)]^1⁄2^, where the first term is the empirical variance of the mean PA across *k* replications and the second term is the sampling variance of a Pearson correlation coefficient for *N* = *N_t_* + *N_p_* genotypes. Finally, we compared the GBLUP approach described above (i.e., based on the GRM alone) to models that also included the first five PC scores of the genomic relationship matrix as fixed-effect covariates (CHEN *et al*. 2015).

We performed GWAS analyses on the *G*(*SR*) and *G* phenotypes using the methods described in EVANS *et al*. (2014) and SLAVOV *et al*. (2014). Briefly, we used the EMMAX software to implement the Efficient Mixed Model Association Expedited approach (KANG *et al*. 2010). Models for all GWAS analyses included the identity-by-state kinship matrix to control for cryptic relatedness and population structure. A subset of analyses also included the first five PCs from the SMARTPCA analyses described above as fixed-effect covariates (PRICE *et al*. 2010).

### Linkage disequilibrium and haplotype sharing

Across all hierarchical levels, we calculated *r*^2^ for each pair of SNPs located within 10 kb of each other using different MAF cut-offs or bins (*SI Appendix*, Materials and Methods, Table S5). We used these data to estimate the probability of tagging a randomly assigned (hypothetical) QTN. This probability was calculated as the proportion of times at least one SNP within 10 kb had *r*^2^ ≥ 0.6 with the hypothetical QTN. For the within-river analyses, we used the same approach but equalized sample sizes within rivers using random subsampling (*SI Appendix*, Materials and Methods).

To quantify linkage phase consistency among rivers, stands-within-rivers, and genotypes-within-stands, we calculated haplotype sharing (DE ROOS *et al*. 2008; GIBBS *et al*. 2009; KIJAS *et al*. 2012) at each of these levels (*SI Appendix*, Materials and Methods).

### Geographic and climatic random forest and ridge regression analyses

We used rrBLUP and random cross-validations to compare the predictive abilities of SNPs versus those obtained using geographic and climatic variables. The geographic variables consisted of latitude, longitude, and elevation, whereas the climate variables consisted of 21 temperature and precipitation related variables from ClimateNA v5.21 (WANG *et al*. 2016).

We used rrBLUP to be consistent with the GBLUP analyses described above. We also used lasso regression to evaluate the relative importance of the geographic and climatic variables. Because lasso regression involves variable selection, it is useful for interpreting the relative importance of the regression predictors. The details of these analyses are described in the *SI Appendix,* Materials and Methods.

### Delineation of seed zones

We evaluated the performance of phenotypes, SNPs, climate variables, and geographic variables for delineating seed zones. Although *Populus* species are typically propagated clonally, the term ‘seed zone’ is often used to denote native populations of forest trees with sufficient genetic homogeneity to be treated as a single population for reforestation purposes. We used three methods to delineate the true or target seed zones, and then compared these to ‘reconstructed’ zones delineated using phenotypes or ridge regression predictions (i.e., for SNPs, climate variables, and geographic variables). We delineated ‘true’ zones by (1) assuming they corresponded to the 16 rivers, (2) using K-means clustering to delineate 16 zones based on the phenotypes, and (3) using K-means clustering to delineate 3 zones based on the phenotypes. To compare the true vs reconstructed zone allocations, we calculated cluster purity (MANNING *et al*. 2008), which is the proportion of stands in each reconstructed seed zone that were also in the same true zone (*SI Appendix*, Materials and Methods).

## Acknowledgments and funding sources

This study was funded by the US DOE Bioenergy Science Center (BESC). We acknowledge Luke Evans for his contributions to discussions during the early phases of the study. We also thank Tal Shalev, Steve Strauss and Michael Nagle for their helpful suggestions and comments on earlier versions of this manuscript.

## References

1. Aitken, S. N., and J. B. Bemmels, 2016 Time to get moving: assisted gene flow of forest trees. Evolutionary Applications 9: 271–290.

2. Aitken, S. N., and M. C. Whitlock, 2013 Assisted Gene Flow to Facilitate Local Adaptation to Climate Change, pp. 367-+ in Annual Review of Ecology, Evolution, and Systematics, Vol 44, edited by D. J. Futuyma.

3. Aitken, S. N., S. Yeaman, J. A. Holliday, T. Wang and S. Curtis-McLane, 2008 Adaptation, migration or extirpation: climate change outcomes for tree populations. Evolutionary Applications 1: 95–111.

4. Alberto, F. J., S. N. Aitken, R. Alia, S. C. Gonzalez-Martinez, H. Hanninen et al., 2013 Potential for evolutionary responses to climate change evidence from tree populations. Global Change Biology 19: 1645–1661.

5. Beavis, W. D., 1998 QTL analyses: Power, precision, and accuracy, pp. 145–162 in Molecular dissection of complex traits, edited by A. H. Paterson. CRC Press, Boca Raton, FL.

6. Beavis, W. D., O. S. Smith, D. Grant and R. Fincher, 1994 Identification of quantitative trait loci using a small sample of topcrossed and F4 progeny from maize. Crop Science 34: 882–896.

7. Bernardo, R., 2016 Bandwagons I, too, have known. Theor Appl Genet 129: 2323–2332.

8. Bhatia, G., N. J. Patterson, S. Sankararaman and A. L. Price, 2013 Estimating and interpreting Fst: the impact of rare variants. Genome Research.

9. Borrell, J. S., J. Zohren, R. A. Nichols and R. J. A. Buggs, 2020 Genomic assessment of local adaptation in dwarf birch to inform assisted gene flow. Evolutionary Applications 13: 161–175.

10. Brown, G. R., D. L. Bassoni, G. P. Gill, J. R. Fontana, N. C. Wheeler et al., 2003 Identification of quantitative trait loci influencing wood property traits in loblolly pine (L.).: III.: QTL verification and candidate gene mapping. Genetics 164: 1537–1546.

11. Butler, J. B., J. S. Freeman, B. M. Potts, R. E. Vaillancourt, H. V. Kahrood et al., 2022 Patterns of genomic diversity and linkage disequilibrium across the disjunct range of the Australian forest tree Eucalyptus globulus. Tree Genetics & Genomes 18: 28.

12. Campbell, R. K., 1979 Genecology of Douglas-Fir in A Watershed in the Oregon Cascades. Ecology 60: 1036–1050.

13. Campbell, R. K., 1986 Mapped Genetic-Variation of Douglas-Fir to Guide Seed Transfer in Southwest Oregon. Silvae Genetica 35: 85–96.

14. Chang, C. C., C. C. Chow, L. C. Tellier, S. Vattikuti, S. M. Purcell et al., 2015 Second-generation PLINK: rising to the challenge of larger and richer datasets. Gigascience 4: 7.

15. Chen, C. Y., J. Han, D. J. Hunter, P. Kraft and A. L. Price, 2015 Explicit Modeling of Ancestry Improves Polygenic Risk Scores and BLUP Prediction. Genet Epidemiol 39: 427–438.

16. Chen, Z. Q., L. Grossfurthner, J. L. Loxterman, J. Masingale, B. A. Richardson et al., 2022 Applying genomics in assisted migration under climate change: Framework, empirical applications, and case studies. Evolutionary Applications 15: 3–21.

17. Chhetri, H. B., A. Furches, D. Macaya-Sanz, A. R. Walker, D. Kainer et al., 2020 Genome-Wide Association Study of Wood Anatomical and Morphological Traits in Populus trichocarpa. Frontiers in Plant Science 11.

18. Curtis, P. G., C. M. Slay, N. L. Harris, A. Tyukavina and M. C. Hansen, 2018 Classifying drivers of global forest loss. Science 361: 1108–1111.

19. Daetwyler, H. D., M. P. L. Calus, R. Pong-Wong, G. de Los Campos and J. M. Hickey, 2013 Genomic prediction in animals and plants: simulation of data, validation, reporting, and benchmarking. Genetics 193: 347–365.

20. Danecek, P., A. Auton, G. Abecasis, C. A. Albers, E. Banks et al., 2011 The variant call format and VCFtools. Bioinformatics 27: 2156–2158.

21. De La Torre, A. R., B. Wilhite, D. Puiu, J. B. St Clair, M. W. Crepeau et al., 2021 Dissecting the Polygenic Basis of Cold Adaptation Using Genome-Wide Association of Traits and Environmental Data in Douglas-fir. Genes (Basel) 12.

22. de Los Campos, G., J. M. Hickey, R. Pong-Wong, H. D. Daetwyler and M. P. L. Calus, 2013 Whole-genome regression and prediction methods applied to plant and animal breeding. Genetics 193: 327–345.

23. de Roos, A. P., B. J. Hayes, R. J. Spelman and M. E. Goddard, 2008 Linkage disequilibrium and persistence of phase in Holstein-Friesian, Jersey and Angus cattle. Genetics 179: 1503–1512.

24. DeBell, D. S., 1990 Populus trichocarpa Torr. & Gray, Black Cottonwood, pp. 570–576 in Silvics of North America Vol. 2. Hardwoods. Agriculture Handbook 654, edited by R. M. Burns and B. H. Honkala. U.S. Department of Agriculture, Forest Service, Washington D.C.

25. Dunlap, J. M., and R. F. Stettler, 1996 Genetic variation and productivity of Populus trichocarpa and its hybrids. IX. Phenology and Melampsora rust incidence of native black cottonwood clones from four river valleys in Washington. Forest Ecology & Management 87: 233–256.

26. Eckert, A. J., A. D. Bower, S. C. Gonzalez-Martinez, J. L. Wegrzyn, G. Coop et al., 2010a Back to nature: ecological genomics of loblolly pine (Pinus taeda, Pinaceae). Molecular Ecology 19: 3789–3805.

27. Eckert, A. J., A. D. Bower, J. L. Wegrzyn, B. Pande, K. D. Jermstad et al., 2009 Asssociation Genetics of Coastal Douglas Fir (Pseudotsuga menziesu var. menziesii, Pinaceae). I. Cold-Hardiness Related Traits. Genetics 182: 1289–1302.

28. Eckert, A. J., J. van Heerwaarden, J. L. Wegrzyn, C. D. Nelson, J. Ross-Ibarra et al., 2010b Patterns of Population Structure and Environmental Associations to Aridity Across the Range of Loblolly Pine (Pinus taeda L., Pinaceae). Genetics 185: 969–982.

29. Endelman, J. B., 2011 Ridge Regression and Other Kernels for Genomic Selection with R Package rrBLUP. Plant Genome 4: 250–255.

30. Evans, L. M., G. T. Slavov, E. Rodgers-Melnick, J. Martin, P. Ranjan et al., 2014 Population genomics of *Populus trichocarpa* identifies signatures of selection and adaptive trait associations. Nat Genet 46: 1089–1096.

31. Fitzpatrick, M. C., V. E. Chhatre, R. Y. Soolanayakanahally and S. R. Keller, 2021 Experimental support for genomic prediction of climate maladaptation using the machine learning approach Gradient Forests. Molecular Ecology Resources 21: 2749–2765.

32. Forde, A., G. Hemani and J. Ferguson, 2023 Review and further developments in statistical corrections for Winner’s Curse in genetic association studies. PLOS Genetics 19: e1010546.

33. Frank, A., G. T. Howe, C. Sperisen, P. Brang, J. B. St Clair et al., 2017 Risk of genetic maladaptation due to climate change in three major European tree species. Global Change Biology 23: 5358–5371.

34. Furches, A., D. Kainer, D. Weighill, A. Large, P. Jones et al., 2019 Finding New Cell Wall Regulatory Genes in Populus trichocarpa Using Multiple Lines of Evidence. Frontiers in Plant Science 10.

35. Gain, C., B. Rhoné, P. Cubry, I. Salazar, F. Forbes et al., 2023 A Quantitative Theory for Genomic Offset Statistics. Molecular Biology and Evolution 40.

36. Garzón, M. B., T. M. Robson and A. Hampe, 2019 ΔTraitSDMs: species distribution models that account for local adaptation and phenotypic plasticity. New Phytologist 222: 1757–1765.

37. Geraldes, A., S. P. Difazio, G. T. Slavov, P. Ranjan, W. Muchero et al., 2013 A 34K SNP genotyping array for Populus trichocarpa: Design, application to the study of natural populations and transferability to other Populus species. Molecular Ecology Resources 13: 306–323.

38. Gibbs, R. A., J. F. Taylor, C. P. Van Tassell, W. Barendse, K. A. Eversole et al., 2009 Genome-wide survey of SNP variation uncovers the genetic structure of cattle breeds. Science 324: 528–532.

39. Gornall, J. L., and R. D. Guy, 2007 Geographic variation in ecophysiological traits of black cottonwood (Populus trichocarpa). Canadian Journal of Botany-Revue Canadienne de Botanique 85: 1202–1213.

40. Goudet, J., 2005 HIERFSTAT, a package for R to compute and test hierarchical F-statistics. Molecular Ecology Notes 5: 184–186.

41. Grattapaglia, D., 2022 Twelve years into genomic selection in forest trees: Climbing the slope of enlightenment of marker assisted breeding. Forests 13, 1554: 1–25.

42. Grattapaglia, D., O. B. Silva-Junior, R. T. Resende, E. P. Cappa, B. S. F. Muller et al., 2018 Quantitative genetics and genomics converge to accelerate forest tree breeding. Front Plant Sci 9: 1693.

43. Gray, L. K., T. Gylander, M. S. Mbogga, P.-y. Chen and A. Hamann, 2011 Assisted migration to address climate change: recommendations for aspen reforestation in western Canada. Ecological Applications 21: 1591–1603.

44. Gray, L. K., and A. Hamann, 2013 Tracking suitable habitat for tree populations under climate change in western North America. Climatic Change 117: 289–303.

45. Hall, D., H. R. Hallingbäck and H. X. Wu, 2016 Estimation of number and size of QTL effects in forest tree traits. Tree Genetics & Genomes 12: 110.

46. Hamann, A., D. R. Roberts, Q. E. Barber, C. Carroll and S. E. Nielsen, 2015 Velocity of climate change algorithms for guiding conservation and management. Global Change Biology 21: 997–1004.

47. Hamrick, J. L., M. J. W. Godt and S. L. Sherman-Broyles, 1992 Factors influencing levels of genetic diversity in woody plant species. New for. 6: 95–124.

48. Hickey, J. M., T. Chiurugwi, I. Mackay and W. Powell, 2017 Genomic prediction unifies animal and plant breeding programs to form platforms for biological discovery. Nat Genet 49: 1297–1303.

49. Hill, A. P., and C. B. Field, 2021 Forest fires and climate-induced tree range shifts in the western US. Nature Communications 12.

50. Hiraoka, Y., E. Fukatsu, K. Mishima, T. Hirao, K. M. Teshima et al., 2018 Potential of Genome-Wide Studies in Unrelated Plus Trees of a Coniferous Species, Cryptomeria japonica (Japanese Cedar). Front Plant Sci 9: 1322.

51. Holliday, J. A., K. Ritland and S. N. Aitken, 2010 Widespread, ecologically relevant genetic markers developed from association mapping of climate-related traits in Sitka spruce (Picea sitchensis). New Phytologist 188: 501–514.

52. Holliday, J. A., T. Wang and S. Aitken, 2012 Predicting Adaptive Phenotypes From Multilocus Genotypes in Sitka Spruce (Picea sitchensis) Using Random Forest. G3- Genes Genomes Genetics 2: 1085–1093.

53. Holliday, J. A., L. C. Zhou, R. Bawa, M. Zhang and R. W. Oubida, 2016 Evidence for extensive parallelism but divergent genomic architecture of adaptation along altitudinal and latitudinal gradients in Populus trichocarpa. New Phytologist 209: 1240–1251.

54. Howe, G. T., S. N. Aitken, D. B. Neale, K. D. Jermstad, N. C. Wheeler et al., 2003 From genotype to phenotype: unraveling the complexities of cold adaptation in forest trees. Canadian Journal of Botany-Revue Canadienne de Botanique 81: 1247–1266.

55. Howe, G. T., D. P. Horvath, P. Dharmawardhana, H. D. Priest, T. C. Mockler et al., 2015 Extensive Transcriptome Changes During Natural Onset and Release of Vegetative Bud Dormancy in Populus. Frontiers in plant science 6: 989–989.

56. Howe, G. T., K. J. Jayawickrama, M. L. Cherry, N. C. Wheeler and G. R. Johnson, 2006 Breeding Douglas-fir, pp. 245-353 in Plant Breeding Reviews, edited by J. Janick. John Wiley and Sons, Inc., Hoboken, New Jersey.

57. Hurel, A., M. de Miguel, C. Dutech, M. L. Desprez-Loustau, C. Plomion et al., 2021 Genetic basis of growth, spring phenology, and susceptibility to biotic stressors in maritime pine. Evol Appl 14: 2750–2772.

58. Ingvarsson, P. K., M. V. Garcia, V. Luquez, D. Hall and S. Jansson, 2008 Nucleotide polymoirphism and phenotypic associations within and around the phytochrome B2 locus in European aspen (Populus tremula, Salicaceae). Genetics 178: 2217–2226.

59. IPCC Core Writing Team, 2023 Climate Change 2023: Synthesis Report. Contribution of Working Groups I, II and III to the Sixth Assessment Report of the Intergovernmental Panel on Climate Change, edited by H. Lee and J. Romero. Intergovernmental Panel on Climate Change, Geneva, Switzerland.

60. Josephs, E. B., J. R. Stinchcombe and S. I. Wright, 2017 What can genome-wide association studies tell us about the evolutionary forces maintaining genetic variation for quantitative traits? New Phytologist 214: 21–33.

61. Kachuri, L., N. Chatterjee, J. Hirbo, D. J. Schaid, I. Martin et al., 2024 Principles and methods for transferring polygenic risk scores across global populations. Nature Reviews Genetics 25: 8–25.

62. Kang, H. M., J. H. Sul, S. K. Service, N. A. Zaitlen, S. Y. Kong et al., 2010 Variance component model to account for sample structure in genome-wide association studies. Nature Genetics 42: 348–U110.

63. Kijas, J. W., J. A. Lenstra, B. Hayes, S. Boitard, L. R. Porto Neto et al., 2012 Genome-Wide Analysis of the World’s Sheep Breeds Reveals High Levels of Historic Mixture and Strong Recent Selection. PLOS Biology 10: e1001258.

64. Kooke, R., W. Kruijer, R. Bours, F. Becker, A. Kuhn et al., 2016 Genome-Wide Association Mapping and Genomic Prediction Elucidate the Genetic Architecture of Morphological Traits in Arabidopsis. Plant Physiology 170: 2187–2203.

65. Langlet, O., 1971 Two Hundred Years Genecology. Taxon 20: 653–721.

66. Láruson, A. J., M. C. Fitzpatrick, S. R. Keller, B. C. Haller and K. E. Lotterhos, 2022 Seeing the forest for the trees: Assessing genetic offset predictions from gradient forest. Evolutionary Applications 15: 403–416.

67. Leites, L., and M. B. Garzon, 2023 Forest tree species adaptation to climate across biomes: Building on the legacy of ecological genetics to anticipate responses to climate change. Global Change Biology.

68. Leites, L. P., A. P. Robinson, G. E. Rehfeldt, J. D. Marshall and N. L. Crookston, 2012 Height-growth response to climatic changes differs among populations of Douglas-fir: a novel analysis of historic data. Ecological Applications 22: 154–165.

69. Lello, L., S. G. Avery, L. Tellier, A. I. Vazquez, G. de los Campos et al., 2018 Accurate Genomic Prediction of Human Height. Genetics 210: 477.

70. Li, Y. R., and B. J. Keating, 2014 Trans-ethnic genome-wide association studies: advantages and challenges of mapping in diverse populations. Genome Medicine 6: 91.

71. Lin, R.-C., B. T. Ferreira and Y.-W. Yuan, 2024 The molecular basis of phenotypic evolution: beyond the usual suspects. Trends in Genetics.

72. Lind, B. M., R. Candido-Ribeiro, P. Singh, M. M. Lu, D. O. Vidakovic et al., 2024 How useful are genomic data for predicting maladaptation to future climate? Global Change Biology 30.

73. Lind, B. M., and K. E. Lotterhos, 2024 The limits of predicting maladaptation to future environments with genomic data. bioRxiv: 2024.2001.2030.577973.

74. Ling, A. S., E. Hay, S. E. Aggrey and R. Rekaya, 2021 Dissection of the impact of prioritized QTL-linked and -unlinked SNP markers on the accuracy of genomic selection. Bmc Genomic Data 22.

75. Lotterhos, K. E., 2019 The Effect of Neutral Recombination Variation on Genome Scans for Selection. G3 (Bethesda) 9: 1851–1867.

76. Lotterhos, K. E., 2023 The paradox of adaptive trait clines with nonclinal patterns in the underlying genes. Proc Natl Acad Sci U S A 120: e2220313120.

77. Mackay, T. F. C., and R. R. H. Anholt, 2024 Pleiotropy, epistasis and the genetic architecture of quantitative traits. Nature Reviews Genetics 25: 639–657.

78. MacLachlan, I. R., T. K. McDonald, B. M. Lind, L. H. Rieseberg, S. Yeaman et al., 2021 Genome-wide shifts in climate-related variation underpin responses to selective breeding in a widespread conifer. Proceedings of the National Academy of Sciences of the United States of America 118.

79. MacLachlan, I. R., T. L. Wang, A. Hamann, P. Smets and S. N. Aitken, 2017 Selective breeding of lodgepole pine increases growth and maintains climatic adaptation. Forest Ecology and Management 391: 404–416.

80. Makowsky, R., N. M. Pajewski, Y. C. Klimentidis, A. I. Vazquez, C. W. Duarte et al., 2011 Beyond Missing Heritability: Prediction of Complex Traits. Plos Genetics 7.

81. Manning, D. C., P. Raghavan and H. Schütze, 2008 Introduction to Information Retrieval. Cambridge University Press, Cambridge, UK.

82. McKown, A. D., R. D. Guy, J. Klapste, A. Geraldes, M. Friedmann et al., 2014a Geographical and environmental gradients shape phenotypic trait variation and genetic structure in Populus trichocarpa. New Phytologist 201: 1263–1276.

83. McKown, A. D., J. Klápště, R. D. Guy, Y. A. El-Kassaby and S. D. Mansfield, 2018 Ecological genomics of variation in bud-break phenology and mechanisms of response to climate warming in Populus trichocarpa. New Phytologist 220: 300–316.

84. McKown, A. D., J. Klapste, R. D. Guy, A. Geraldes, I. Porth et al., 2014b Genome-wide association implicates numerous genes underlying ecological trait variation in natural populations of Populus trichocarpa. New Phytologist 203: 535–553.

85. Meuwissen, T. H., B. J. Hayes and M. E. Goddard, 2001 Prediction of total genetic value using genome-wide dense marker maps. Genetics 157: 1819–1829.

86. Millar, C. I., and R. D. Westfall, 1992 Allozyme markers in forest genetic conservation. New Forests 6: 347–371.

87. Millennium Ecosystem Assessment, 2005 Ecosystems and Human Well-being: Synthesis. World Resources Institute, Island Press, Washington, DC.

88. Morgenstern, E. K., 1996 Geographic variation in forest trees: genetic basis and application of knowledge in silviculture. Univ. British Columbia Press, Vancouver, B.C.

89. Müller, B. S. F., J. E. de Almeida Filho, B. M. Lima, C. C. Garcia, A. Missiaggia et al., 2019 Independent and Joint-GWAS for growth traits in Eucalyptus by assembling genome-wide data for 3373 individuals across four breeding populations. New Phytol 221: 818–833.

90. Müller, B. S. F., L. G. Neves, J. E. de Almeida Filho, M. F. R. Resende, Jr., P. R. Muñoz et al., 2017 Genomic prediction in contrast to a genome-wide association study in explaining heritable variation of complex growth traits in breeding populations of Eucalyptus. BMC genomics 18: 524–524.

91. Neale, D. B., and A. Kremer, 2011 Forest tree genomics: growing resources and applications. Nat Rev Genet 12: 111–122.

92. Neale, D. B., and O. Savolainen, 2004 Association genetics of complex traits in conifers. Trends in Plant Science 9: 325–330.

93. Nelson, M. R., K. Bryc, K. S. King, A. Indap, A. R. Boyko et al., 2008 The population reference sample, POPRES: A resource for population, disease, and pharmacological genetics research. American Journal of Human Genetics 83: 347–358.

94. Park, A., and C. Talbot, 2018 Information Underload: Ecological Complexity, Incomplete Knowledge, and Data Deficits Create Challenges for the Assisted Migration of Forest Trees. Bioscience 68: 251–263.

95. Patterson, N., A. L. Price and D. Reich, 2006 Population structure and eigenanalysis. Plos Genetics 2: 2074–2093.

96. Pauley, S. S., and T. O. Perry, 1954 Ecotypic variation of the photoperiodic response in Populus. Journal of the Arnold Arboretum 35: 167–188.

97. Perez, P., and G. de los Campos, 2014 Genome-Wide Regression and Prediction with the BGLR Statistical Package. Genetics 198: 483–U463.

98. Petit, R. J., and A. Hampe, 2006 Some evolutionary consequences of being a tree, pp. 187-214 in Annual Review of Ecology Evolution and Systematics.

99. Porth, I., J. Klapste, A. D. McKown, J. La Mantia, R. D. Guy et al., 2015 Evolutionary Quantitative Genomics of Populus trichocarpa. Plos One 10.

100. Price, A. L., N. A. Zaitlen, D. Reich and N. Patterson, 2010 New approaches to population stratification in genome-wide association studies. Nature Reviews Genetics 11: 459–463.

101. R Core Team, 2021 R: A language and environment for statistical computing. R Foundation for Statistical Computing, Vienna, Austria, pp.

102. Rehfeldt, G. E., N. M. Tchebakova, Y. I. Parfenova, W. R. Wykoff, N. A. Kuzmina et al., 2002 Intraspecific responses to climate in Pinus sylvestris. Global Change Biology 8: 912–929.

103. Rehfeldt, G. E., C. C. Ying, D. L. Spittlehouse and D. A. Hamilton, 1999 Genetic responses to climate in Pinus contorta: Niche breadth, climate change, and reforestation. Ecological Monographs 69: 375–407.

104. Rellstab, C., B. Dauphin and M. Exposito-Alonso, 2021 Prospects and limitations of genomic offset in conservation management. Evol Appl 14: 1202–1212.

105. Rohde, A., T. Ruttink, V. Hostyn, L. Sterck, K. Van Driessche et al., 2007 Gene expression during the induction, maintenance, and release of dormancy in apical buds of poplar. Journal of Experimental Botany 58: 4047–4060.

106. Scotti, I., S. C. Gonzalez-Martinez, K. B. Budde and H. Lalague, 2016 Fifty years of genetic studies: what to make of the large amounts of variation found within populations? Annals of Forest Science 73: 69–75.

107. Shalev, T. J., O. Gamal El-Dien, M. M. S. Yuen, S. Shengqiang, S. D. Jackman et al., 2022 The western redcedar genome reveals low genetic diversity in a self-compatible conifer. Genome Research 32: 1952–1964.

108. Silva-Junior, O. B., and D. Grattapaglia, 2015 Genome-wide patterns of recombination, linkage disequilibrium and nucleotide diversity from pooled resequencing and single nucleotide polymorphism genotyping unlock the evolutionary history of Eucalyptus grandis. New Phytologist 208: 830–845.

109. Slavov, G. T., S. P. DiFazio, J. Martin, W. Schackwitz, W. Muchero et al., 2012 Genome resequencing reveals multiscale geographic structure and extensive linkage disequilibrium in the forest tree Populus trichocarpa. New Phytologist 196: 713–725.

110. Slavov, G. T., S. P. DiFazio and S. H. Strauss, 2004 Gene flow in forest trees: Gene migration patterns and landscape modeling of transgene dispersion in hybrid poplar, pp. 89–106 in Introgression from Genetically Modified Plants into Wild Relatives, edited by J. C. M. den Nijs, D. Bartsch and J. Sweet. CAB International, UK.

111. Slavov, G. T., R. Nipper, P. Robson, K. Farrar, G. G. Allison et al., 2014 Genome-wide association studies and prediction of 17 traits related to phenology, biomass and cell wall composition in the energy grass Miscanthus sinensis. New Phytologist 201: 1227–1239.

112. Slavov, G. T., and P. Zhelev, 2010 Salient biological features, systematics, and genetic variation of Populus pp. 15-38 in Genetics and genomics of Populus edited by S. Jansson, R. Bhalerao and A. T. Groover. Springer, NY.

113. St Clair, J. B., and G. T. Howe, 2007 Genetic maladaptation of coastal Douglas-fir seedlings to future climates. Global Change Biology 13: 1441–1454.

114. St Clair, J. B., N. L. Mandel and K. W. Vance-Borland, 2005 Genecology of Douglas fir in western Oregon and Washington. Annals of Botany 96: 1199–1214.

115. St. Clair, J. B., G. T. Howe and J. G. Kling, 2019 The 1912 Douglas-Fir Heredity Study: Long-Term Effects of Climatic Transfer Distance on Growth and Survival. Journal of Forestry 118: 1–13.

116. Strauss, S. H., G. T. Slavov and S. P. DiFazio, 2022 Gene-editing for production traits in forest trees: Challenges to integration and gene target identification. Forests 13: 1–13.

117. Tan, B. Y., and P. K. Ingvarsson, 2022 Integrating genome-wide association mapping of additive and dominance genetic effects to improve genomic prediction accuracy in *Eucalyptus*. Plant Genome 15.

118. Timpson, N. J., C. M. T. Greenwood, N. Soranzo, D. J. Lawson and J. B. Richards, 2018 Genetic architecture: the shape of the genetic contribution to human traits and disease. Nature Reviews Genetics 19: 110–124.

119. Tuskan, G. A., S. DiFazio, S. Jansson, J. Bohlmann, I. Grigoriev et al., 2006 The Genome of Black Cottonwood, Populus trichocarpa (Torr. & Gray). Science 313: 1596–1604.

120. Visscher, Peter M., Matthew A. Brown, Mark I. McCarthy and J. Yang, 2012 Five Years of GWAS Discovery. The American Journal of Human Genetics 90: 7–24.

121. Visscher, P. M., N. R. Wray, Q. Zhang, P. Sklar, M. I. McCarthy et al., 2017 10 Years of GWAS Discovery: Biology, Function, and Translation. The American Journal of Human Genetics 101: 5–22.

122. Wadgymar, S. M., M. L. DeMarche, E. B. Josephs, S. N. Sheth and J. T. Anderson, 2022 Local Adaptation: Causal Agents of Selection and Adaptive Trait Divergence. Annual Review of Ecology Evolution and Systematics 53: 87–111.

123. Wang, J., J. H. Ding, B. Y. Tan, K. M. Robinson, I. H. Michelson et al., 2018 A major locus controls local adaptation and adaptive life history variation in a perennial plant. Genome Biology 19.

124. Wang, T., A. Hamann, A. Yanchuk, G. A. O’Neill and S. N. Aitken, 2006 Use of response functions in selecting lodgepole pine populations for future climates. Global Change Biology 12: 2404–2416.

125. Wang, T., G. A. O’Neill and S. N. Aitken, 2010 Integrating environmental and genetic effects to predict responses of tree populations to climate. Ecological Applications 20: 153–163.

126. Wang, T. L., A. Hamann, D. Spittlehouse and C. Carroll, 2016 Locally Downscaled and Spatially Customizable Climate Data for Historical and Future Periods for North America. Plos One 11.

127. Wang, Y., J. Guo, G. Ni, J. Yang, P. M. Visscher et al., 2020 Theoretical and empirical quantification of the accuracy of polygenic scores in ancestry divergent populations. Nature Communications 11: 3865.

128. Weber, J. C., R. F. Stettler and P. E. Heilman, 1985 GENETIC-VARIATION AND PRODUCTIVITY OF POPULUS-TRICHOCARPA AND ITS HYBRIDS .1. MORPHOLOGY AND PHENOLOGY OF 50 NATIVE CLONES. Canadian Journal of Forest Research-Revue Canadienne De Recherche Forestiere 15: 376–383.

129. Weiss, M., R. A. Sniezko, D. Puiu, M. W. Crepeau, K. Stevens et al., 2020 Genomic basis of white pine blister rust quantitative disease resistance and its relationship with qualitative resistance. Plant J 104: 365–376.

130. Westfall, R. D., and M. T. Conkle, 1992 Allozyme markers in breeding zone designation. New Forests 6: 279–309.

131. Wray, N. R., 2005 Allele frequencies and the r2 measure of linkage disequilibrium: impact on design and interpretation of association studies. Twin Res Hum Genet 8: 87–94.

132. Yang, J., S. H. Lee, M. E. Goddard and P. M. Visscher, 2011 GCTA: A Tool for Genome-wide Complex Trait Analysis. American Journal of Human Genetics 88: 76–82.

133. Ye, Z. Y., G. A. O’Neill and T. L. Wang, 2022 Climate data for field trials: onsite micro stations versus ClimateNA. Canadian Journal of Forest Research 52: 1028–1041.

134. Yengo, L., S. Vedantam, E. Marouli, J. Sidorenko, E. Bartell et al., 2022 A saturated map of common genetic variants associated with human height. Nature 610: 704-+.

