## Supporting Information for "Population structure limits inferences from genomic prediction and genome-wide association studies in a forest tree"

### Supporting Results

*Predictive ability increased with training population size and number of markers*

As expected, GBLUP PAs increased with the size of the training population (*N*) and when the number of markers (*M*) was increased from 10 to 1,000,000 (Fig. S1). These trends differed between analyses using phenotypic variation across all hierarchical levels (Fig. S1A, B) and those for genotypes within stands (Fig. S1C, D). Predictive abilities tended to increase with *N* at a higher rate in the within-stand analyses than in analyses conducted across all hierarchical levels (Fig. S1A, C). Furthermore, while increasing *M* beyond 100,000 had little effect on PA in analyses across all hierarchical levels, a plateau was reached at a higher *M* (1,000,000) in analyses within stands (Fig. S1B, D). In contrast, SNP filtering had little effect on GBLUP PAs. For example, the mean PA across traits was the same (0.735) using SNPs filtered with strict versus liberal filtering criteria (Tables S3 and S4). Furthermore, PA was only slightly lower (0.702) using a much smaller set of 51,820 simulated RAD-Seq markers (Tables S3 and S4).

*Genotype × environment interactions had little effect of predictive abilities*

We examined the magnitude of genotype-by-environment (G×E) interactions using variance components from analyses of phenotypic data and among-plantation genetic correlations for each trait. GxE variance accounted for only 9-20% of the total genetic variation, and genetic correlations ranged from 0.80 to 0.95 (Fig. S5A). Consequently, there was only a small decrease in PA when GBLUP models of phenotypic data from one test plantation were trained on phenotypic data from another (Fig. S5B).

### Supporting Tables and Figures

**Table S1. Stand-level prediction of adaptive traits and SNP PC scores using random forest (RF) and ridge regression (RR).** Dependent variables were stand-level BLUPs for quantitative traits (bud flush, BF; bud set, BS; and height growth, HT) and stand-level PC scores for single-nucleotide polymorphisms (SNPs). Predictive abilities (PA)* were calculated using RF and RR and four sets of independent variables: (1) subsets of climate variables selected using RF at the stand level, (2) all 21 climate variables, (3) geographic variables (latitude, longitude, and elevation), and (4) all climate and geographic variables.[†] SE is the mean standard error of PA across all models and analyses (RF and RR) for each trait. Each SE was based on 100 model runs.

| | | RF-selected climate variables | | All climate variables | | Geographic variables | | All climate and geographic variables | | |
|---|---|---|---|---|---|---|---|---|---|---|
| Trait | Climate variables selected using random forest at the stand-level | RF | RR | RF | RR | RF | RR | RF | RR | SE |
| ***Quantitative traits*** | | | | ***Predictive ability*** | | | | | | |
| BF | Eref + EXT + DD<0 + TD + MCMT + FFP | 0.851 | 0.791 | 0.831 | 0.793 | 0.899 | 0.746 | 0.877 | 0.808 | 0.0010 |
| BS | Eref + DD<0 + TD | 0.919 | 0.866 | 0.900 | 0.903 | 0.976 | 0.950 | 0.976 | 0.965 | 0.0004 |
| HT | Eref + DD<0 + TD | 0.910 | 0.839 | 0.885 | 0.889 | 0.951 | 0.880 | 0.949 | 0.922 | 0.0005 |
| ***SNP PC scores*** | | | | | | | | | | |
| SPC1 | Eref + DD<0 + MCMT + TD + PAS + MSP + SHM | 0.920 | 0.654 | 0.900 | 0.730 | 0.931 | 0.739 | 0.931 | 0.818 | 0.0012 |
| SPC2 | Eref | 0.818 | -0.089 | 0.595 | 0.107 | 0.822 | -0.028 | 0.728 | 0.205 | 0.0140 |
| SPC3 | Eref + TD + CMD + EMT + MSP + bFFP | 0.760 | 0.656 | 0.729 | 0.689 | 0.913 | 0.666 | 0.891 | 0.756 | 0.0015 |
| SPC4 | MSP + SHM + CMD + DD>5 + Eref + DD<18 + DD<0 + EMT + TD | 0.849 | 0.733 | 0.826 | 0.774 | 0.956 | 0.791 | 0.938 | 0.873 | 0.0008 |
| SPC5 | Eref | 0.534 | -0.032 | 0.334 | -0.011 | 0.469 | 0.011 | 0.373 | -0.010 | 0.0087 |

[*] Predictive ability is the correlation between the input dependent values (BLUPs or PC scores) and the values predicted using RF or RR using 10-fold cross-validation.

[†] Annual climate variables are the day of the year on which the frost-free period begins (bFFP), Hargreaves climatic moisture deficit (CMD), degree-days below 0°C (DD<0), degree-days below 18°C (DD<18), degree-days above 5°C (DD>5), extreme minimum temperature over 30 years (EMT), Hargreaves reference evaporation (Eref), extreme maximum temperature over 30 years (EXT), frost-free period (FFP), mean coldest month temperature (MCMT), May to September precipitation (MSP), mean warmest month temperature (MWMT), summer heat-moisture index (SHM = MWMT/(MSP/1000)), precipitation as snow (PAS), and temperature difference between MWMT and MCMT (TD)(WANG *et al.* 2016).

**Table S2. River-level prediction of adaptive traits and SNP PC scores using random forest (RF) and ridge regression (RR).** Dependent variables were river-level BLUPS for quantitative traits (bud flush, BF; bud set, BS; and height growth, HT) and river-level PC scores for SNPs. Predictive abilities (PA)* were calculated using RF and RR and 4 sets of independent variables: (1) subsets of climate variables selected using RF at the site-level, (2) all 21 climate variables, (3) geographic variables (latitude, longitude, and elevation), and (4) all climate and geographic variables.[†] SE is the mean standard error of PA across all models and analyses (RF and RR) for each trait. Each SE was based on 100 model runs.

| | | RF-selected climate variables | | All climate variables | | Geographic variables | | All climate and geographic variables | | |
|---|---|---|---|---|---|---|---|---|---|---|
| Trait | Climate variables selected using random forest at the site-level | RF | RR | RF | RR | RF | RR | RF | RR | SE |
| ***Quantitative traits*** | | | | | | | | | | |
| BF | Eref + EXT + DD<0 + TD + MCMT + FFP | 0.753 | 0.865 | 0.440 | 0.338 | 0.467 | 0.100 | 0.410 | 0.347 | 0.0071 |
| BS | Eref + DD<0 + TD | 0.848 | 0.888 | 0.853 | 0.775 | 0.850 | 0.908 | 0.874 | 0.804 | 0.0028 |
| HT | Eref + DD<0 + TD | 0.830 | 0.853 | 0.902 | 0.850 | 0.900 | 0.875 | 0.908 | 0.869 | 0.0033 |
| ***SNP PC scores*** | | | | | | | | | | |
| PC1 | Eref + DD<0 + MCMT + TD + PAS + MSP + SHM | 0.423 | -0.230 | 0.401 | -0.177 | 0.456 | -0.008 | 0.404 | -0.181 | 0.0145 |
| PC2 | Eref | -0.830 | -0.693 | -0.421 | -0.703 | -0.449 | -0.651 | -0.434 | -0.702 | 0.0123 |
| PC3 | Eref + TD + CMD + EMT + MSP + bFFP | 0.553 | 0.370 | 0.515 | 0.486 | 0.592 | 0.396 | 0.523 | 0.498 | 0.0037 |
| PC4 | MSP + SHM + CMD + DD>5 + Eref + DD<18 + DD<0 + EMT + TD | 0.751 | 0.577 | 0.773 | 0.674 | 0.823 | 0.580 | 0.788 | 0.686 | 0.0023 |
| PC5 | Eref | 0.111 | -0.555 | 0.020 | -0.570 | -0.322 | -0.555 | 0.005 | -0.586 | 0.0083 |

[*] Predictive ability is the correlation between the input dependent values (BLUPS or PC scores) and the values predicted using RF or RR using 8-fold cross-validation.

[†] Annual climate variables are the day of the year on which the frost-free period begins (bFFP), Hargreaves climatic moisture deficit (CMD), degree-days below 0°C (DD<0), degree-days below 18°C (DD<18), degree-days above 5°C (DD>5), extreme minimum temperature over 30 years (EMT), Hargreaves reference evaporation (Eref), extreme maximum temperature over 30 years (EXT), frost-free period (FFP), mean coldest month temperature (MCMT), May to September precipitation (MSP), mean warmest month temperature (MWMT), summer heat-moisture index (SHM = MWMT/(MSP/1000)), precipitation as snow (PAS), and temperature difference between MWMT and MCMT (TD) (Wang et al, 2016).

**Table S3. Predictive abilities for phenotypic values across all hierarchical levels (genotypes, stands, and rivers) using GBLUP (SNP predictor variables) or ridge regression (geographic or climate variables) for 840 *P. trichocarpa* clonal genotypes.** SNP (single-nucleotide polymorphism) markers were filtered using strict, liberal, or '*Rad-Seq*' marker (Rad) criteria (Table S4). All predictive abilities are based on 100 random, 10-fold cross-validations with a training population size of 756 and a prediction population size of 84. Numbers in parentheses are standard errors (see Materials and Methods in main text).

| | | Predictive ability (PA) | | | | |
|---|---|---|---|---|---|---|
| | | GBLUP* | | | Ridge regression (RR)† | |
| Trait‡ | $H^{2}$§ | Strict SNP markers | Liberal SNP markers | Rad markers | Geographic variables | Climate variables |
| BF | 0.794 | 0.631 (0.027) | 0.635 (0.027) | 0.598 (0.028) | 0.531 (0.029) | 0.579 (0.029) |
| BS | 0.512 | 0.784 (0.022) | 0.780 (0.022) | 0.740 (0.024) | 0.731 (0.024) | 0.728 (0.024) |
| HT | 0.427 | 0.789 (0.021) | 0.791 (0.021) | 0.767 (0.023) | 0.714 (0.025) | 0.742 (0.025) |
| ***Average*¶** | **0.578** | **0.735** | **0.735** | **0.702** | **0.659** | **0.683** |

*GBLUP is the predictive ability using 3,175,407 markers filtered using 'strict' criteria, 20,770,783 markers filtered using 'liberal' criteria, or 51,820 markers filtered using '*RAD-Seq*' criteria (Table S4).

† Ridge regression is the predictive ability using ridge regression with 3 geographic variables or 21 climate variables.

‡ Trait is the BLUP prediction for bud flush (BF), bud set (BS), or height growth (HT).

§ $H^2$ is the broad-sense heritability.

¶ Average is the overall average across traits.

**Table S4. Filtering criteria for single-nucleotide polymorphism (SNP) data from 840 *Populus trichocarpa* genotypes.**

| **Filtering criteria**[*] | ***Strict*** | ***Liberal*** | ***RAD-Seq*** |
|---|---|---|---|
| *Min depth* | 8 | NA | NA |
| *GATK genotype quality* | 20 | NA | NA |
| *Missing (%)* | 10 | NA | NA |
| *Minor allele count* | 17[†] | 2 | 2 |
| *Minor allele frequency* | 0.01 | 0.0012[‡] | 0.0012[‡] |
| *No. of alleles* | 2 | 2 | 2 |
| **Statistics**[§] | ***Strict*** | ***Liberal*** | ***RAD-Seq*** |
| *No. of loci* | 3,175,407 | 20,770,783 | 51,820 |
| *Max gap (bp)* | 702,736 | 56,822 | 2,033,113 |
| *No. >10 kb gaps* | 2,112 | 355 | 7,605 |
| *Ave gap (bp)* | 124 | 20 | 7,576 |
| *Med gap (bp)* | 29 | 9 | 36 |
| *Ave missing (%)* | 6.6 | 11.0 | 1.4 |
| *Ave MAF* | 0.15 | 0.08 | 0.09 |
| *Med MAF* | 0.10 | 0.02 | 0.03 |

[*] Filtering criteria include *Min depth*: minimum number of reads per genotype; *GATK genotype quality*: Phred-scaled confidence that the genotype assignment is correct (i.e. difference between the log10-transformed likelihoods of the second most likely and the most likely genotype); *Missing (%)*: maximum percentage of missing genotype data allowed for a given locus; *Minor allele count*: minimum number of copies of the minor allele among all genotypes; *Minor allele frequency*: minimum frequency of the minor allele among all genotypes; and *No. of alleles*: number of SNP alleles detected; and NA: not applicable.

[†] Calculated from minor allele frequency.

[‡] Calculated from minor allele count.

[§] *Statistics include No. of loci*: number of SNPs that passed all filtering criteria; *Max gap (bp)*: largest distance between adjacent SNPs; *No. >10 kb gaps*: number of distances between adjacent SNPs exceeding 10 kb; *Ave gap (bp)*: average distance between adjacent SNPs; *Med gap (bp)*: median distance between adjacent SNPs; *Ave missing (%)*: average percentage of missing genotype data; *Ave MAF*: average minor allele frequency; *Ave MAF*: median minor allele frequency.

**Table S5. Minor allele frequency (MAF) bins[*] used for analyses of linkage disequilibrium.**

| Bin | $MAF_{min}$ | $MAF_{max}$ | $MAF_{midpoint}$ | $N_{SNPs}$ | Ave. gap (bp) | Median gap (bp) |
|---|---|---|---|---|---|---|
| 1 | 0.0012 | 0.0024 | 0.0018 | 2,790,927 | 146 | 62 |
| 2 | 0.0024 | 0.0047 | 0.0036 | 2,482,036 | 166 | 65 |
| 3 | 0.0047 | 0.0094 | 0.0071 | 2,335,579 | 177 | 65 |
| 4 | 0.0094 | 0.0187 | 0.0141 | 2,204,361 | 187 | 68 |
| 5 | 0.0187 | 0.0367 | 0.0277 | 2,096,945 | 195 | 69 |
| 6 | 0.0367 | 0.0709 | 0.0538 | 2,123,088 | 192 | 68 |
| 7 | 0.0709 | 0.1324 | 0.1016 | 2,075,169 | 195 | 67 |
| 8 | 0.1324 | 0.2338 | 0.1831 | 1,870,724 | 216 | 68 |
| 9 | 0.2338 | 0.3790 | 0.3064 | 1,664,173 | 242 | 71 |
| 10 | 0.3790 | 0.5000 | 0.4395 | 1,127,781 | 355 | 83 |

[*] $MAF_{min}$ and $MAF_{max}$ were the lower and upper bounds of bins calculated to assure that all pairs of loci in a bin could have an $r^2$ with a theoretical maximum of at least 0.5 (WRAY 2005); $MAF_{midpoint}$ were the midpoints of each bin [($MAF_{min}$ + $MAF_{max}$)/2]; $N_{SNPs}$ were the number of SNPs in each bin; Ave. gap (bp) were the average gap sizes (i.e., distance between adjacent SNPs) and Median gap (bp) were the median gap sizes for each bin.

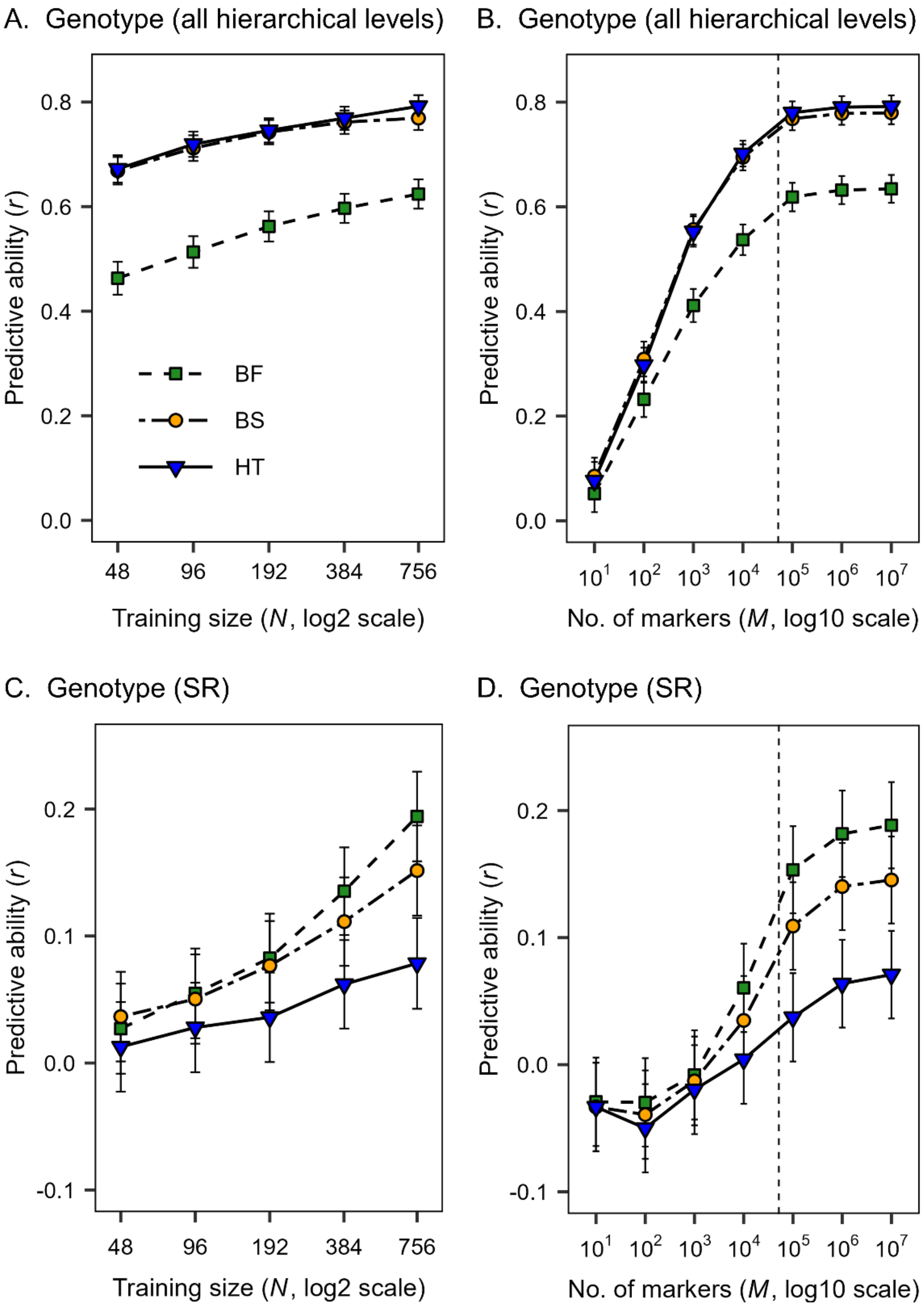


**Figure S1. Predictive ability versus training population size (A, C) and number of markers (B, D) using 840 clonal genotypes.** GBLUP analyses were conducted across all hierarchical levels (A, B) and for genotypes-within-stands-and-rivers [Genotype (SR)] (C, D) using single-nucleotide polymorphism (SNP) markers filtered using the 'liberal' criteria (Table S4). In A and C, training population size ranged from 48 to 756, with a fixed prediction population size of 84, and 20,770,783 SNPs. In B and D, the number of SNPs ranged from 10 to 10,000,000, using a training population size of 756 and a prediction population size of 84. Averages were based on 100 replications and standard errors (error bars) were calculated as described in the main text (Materials and Methods). The dashed line shows the number of RAD-Seq markers.

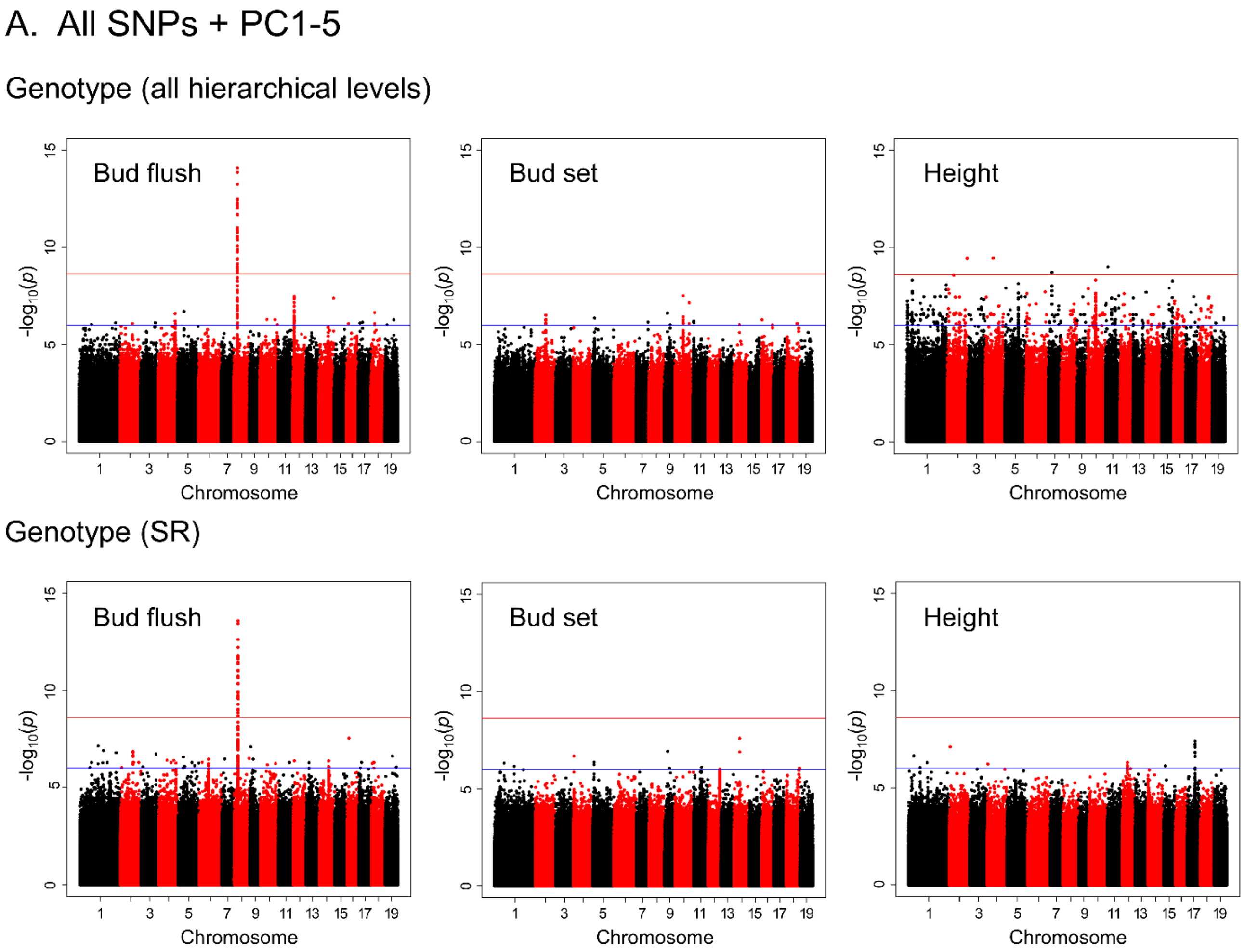


**Figure S2. Genome-wide association study (GWAS) results.** Results are for vegetative bud flush, bud set, and height based on total genetic variation across rivers, stands, and genotypes (A, B, and C top panels) and among genotypes within stands (A, B, and C bottom panels). (A) Manhattan plots from GWAS based on 20,770,783 SNPs filtered using the 'liberal' criteria (Table S4). The first five SNP principal components (SPC1-SPC5) were used to correct for population structure. (B) Same as for panel A except analyses excluded SNPs with MAF < 0.01. (C) Same as for panel A except no SNP principal components were used to correct for population structure and analyses were restricted to 490 genotypes from the Skagit, Puyallup, and Columbia Rivers.

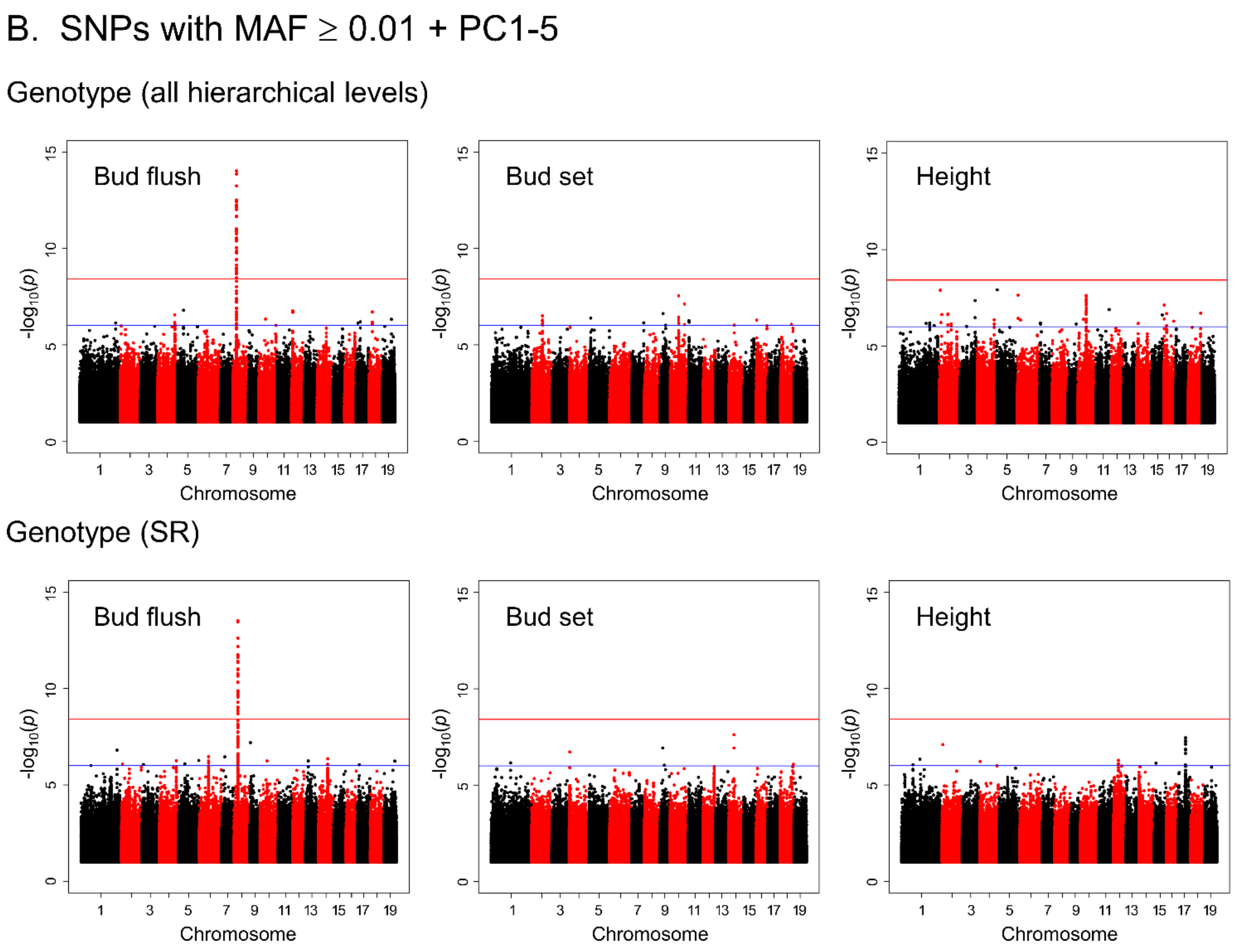


**Figure S2 (cont). Genome-wide association study (GWAS) results.** Results are for vegetative bud flush, bud set, and height based on total genetic variation across rivers, stands, and genotypes (A, B, and C top panels) and among genotypes within stands (A, B, and C bottom panels). (A) Manhattan plots from GWAS based on all 20,770,783 SNPs filtered using the 'liberal' criteria (Table S4). The first five SNP principal components (SPC1-SPC5) were used to correct for population structure. (B) Same as for panel A except analyses excluded SNPs with MAF < 0.01. (C) Same as for panel A except no SNP principal components were used to correct for population structure and analyses were restricted to 490 genotypes from the Skagit, Puyallup, and Columbia Rivers.

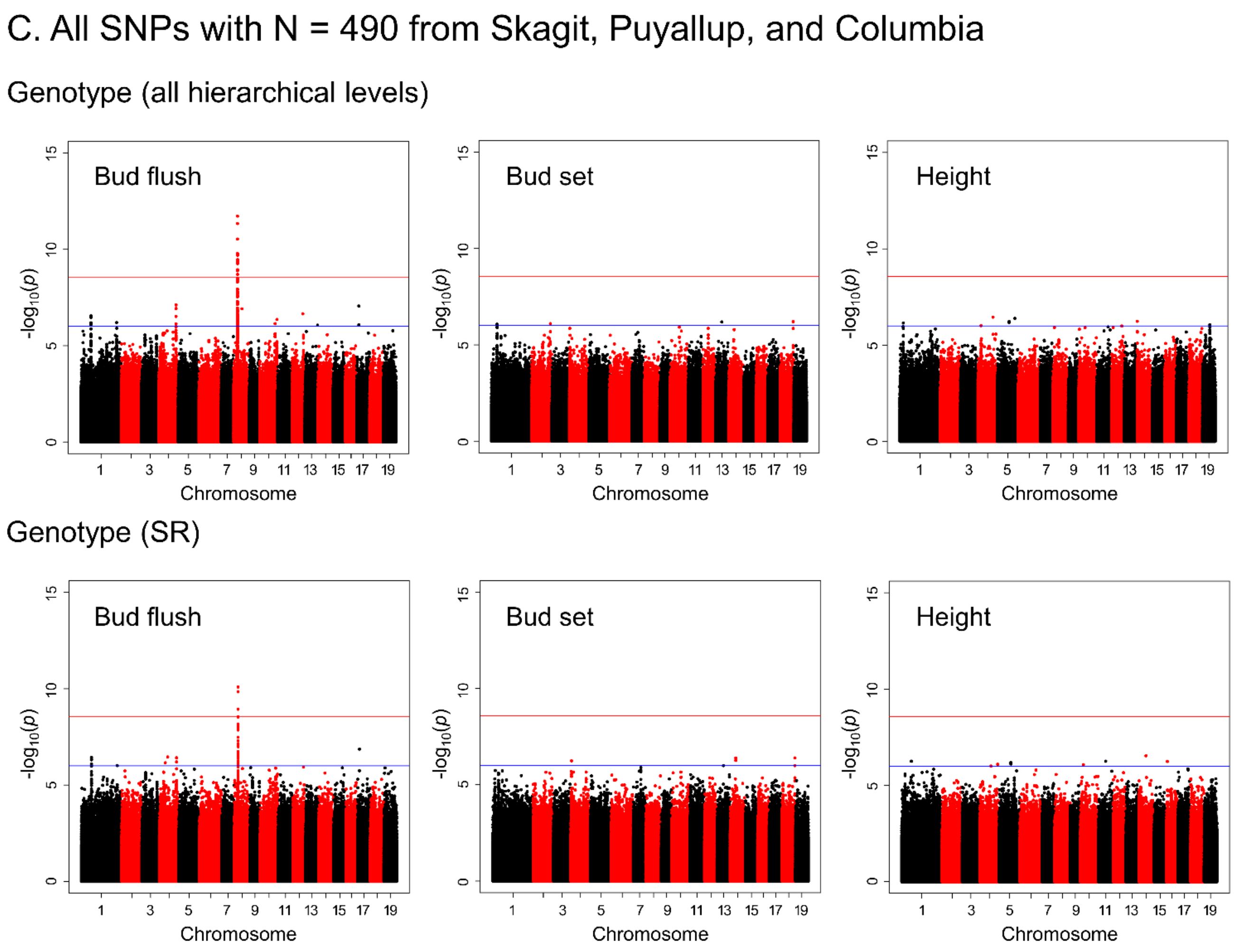


**Figure S2 (cont). Genome-wide association study (GWAS) results.** Results are for vegetative bud flush, bud set, and height based on total genetic variation across rivers, stands, and genotypes (A, B, and C top panels) and among genotypes within stands (A, B, and C bottom panels). (A) Manhattan plots from GWAS based on all 20,770,783 SNPs filtered using the 'liberal' criteria (Table S4). The first five SNP principal components (SPC1-SPC5) were used to correct for population structure. (B) Same as for panel A except analyses excluded SNPs with MAF $< 0.01$. (C) Same as for panel A except no SNP principal components were used to correct for population structure and analyses were restricted to 490 genotypes from the Skagit, Puyallup, and Columbia Rivers.

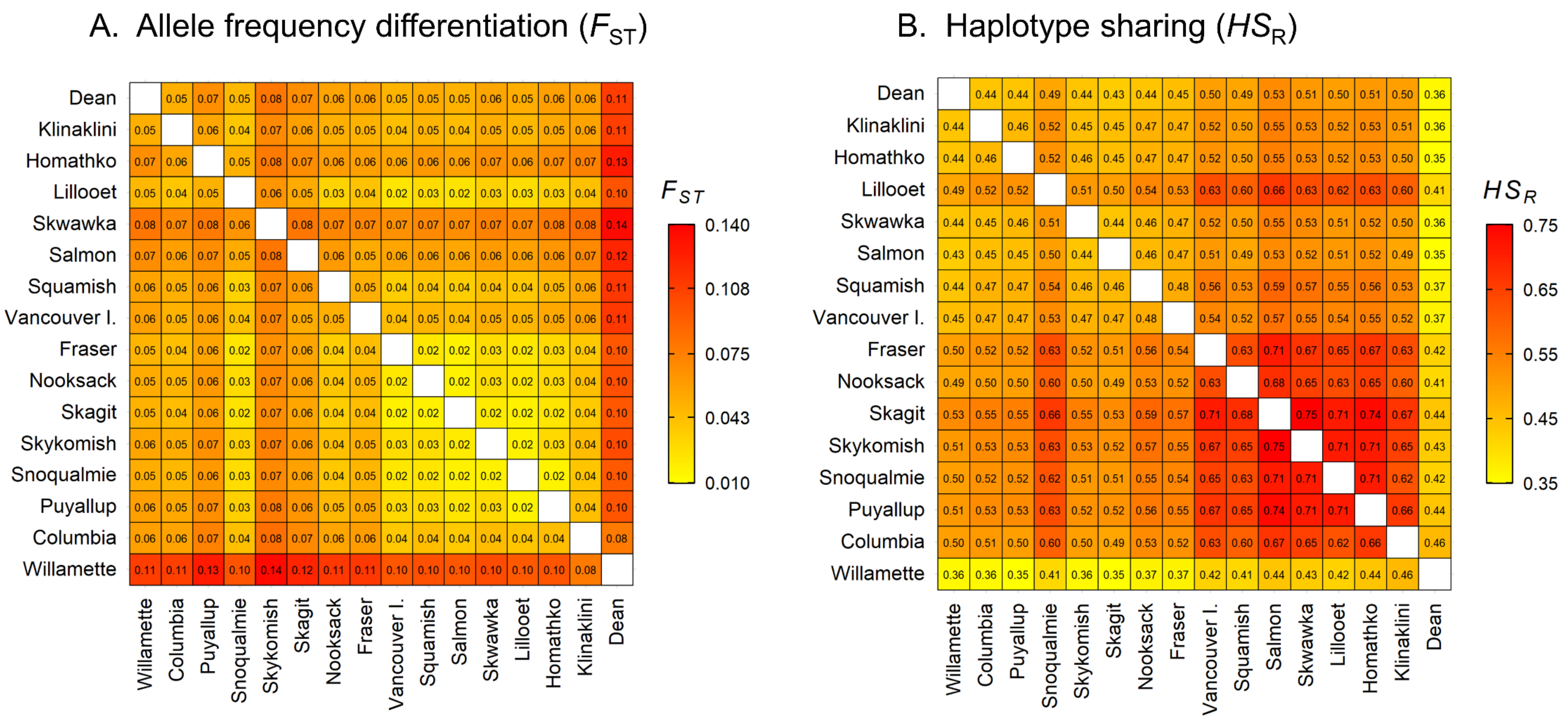


**Figure S3. Allele frequency differentiation ($F_{ST}$) and linkage phase consistency (haplotype sharing) among rivers.** (A) Pairwise $F_{ST}$ values (Hudson's estimator) were calculated at the river level using SMARTPCA. (B) Haplotype sharing analyses were based on 1,082,633 SNPs with MAF ≥ 0.01 separated by at least 300 bp and haplotype sharing was calculated for all pairs of SNPs located within 10 kb of each other.

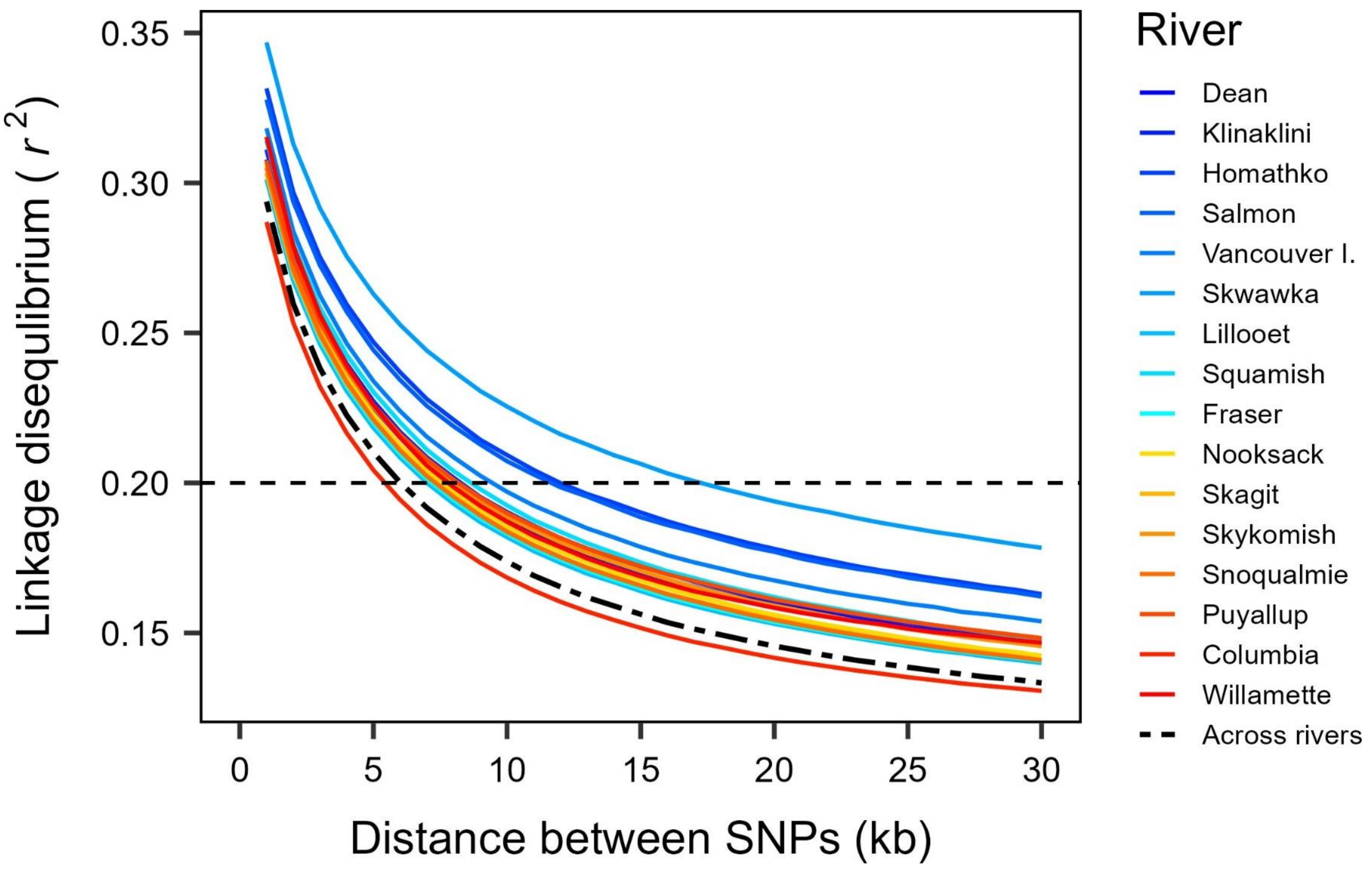


**Figure S4. Linkage disequilibrium (LD) within and across rivers.** LD was calculated as the average $r^2$ for pairs of SNPs with MAF ≥ 0.10 in each 1-kb distance class. To equalize sample sizes, $N$ = 10 clonal genotypes were randomly sampled from each river and 10 were sampled across rivers (i.e., one from each of 10 randomly sampled rivers). Average $r^2$ values were calculated based on 10 random sampling replications and corrected for the small sample size by subtracting $1/2N$. Because of the large number of SNPs, the standard error of each data point was near zero.

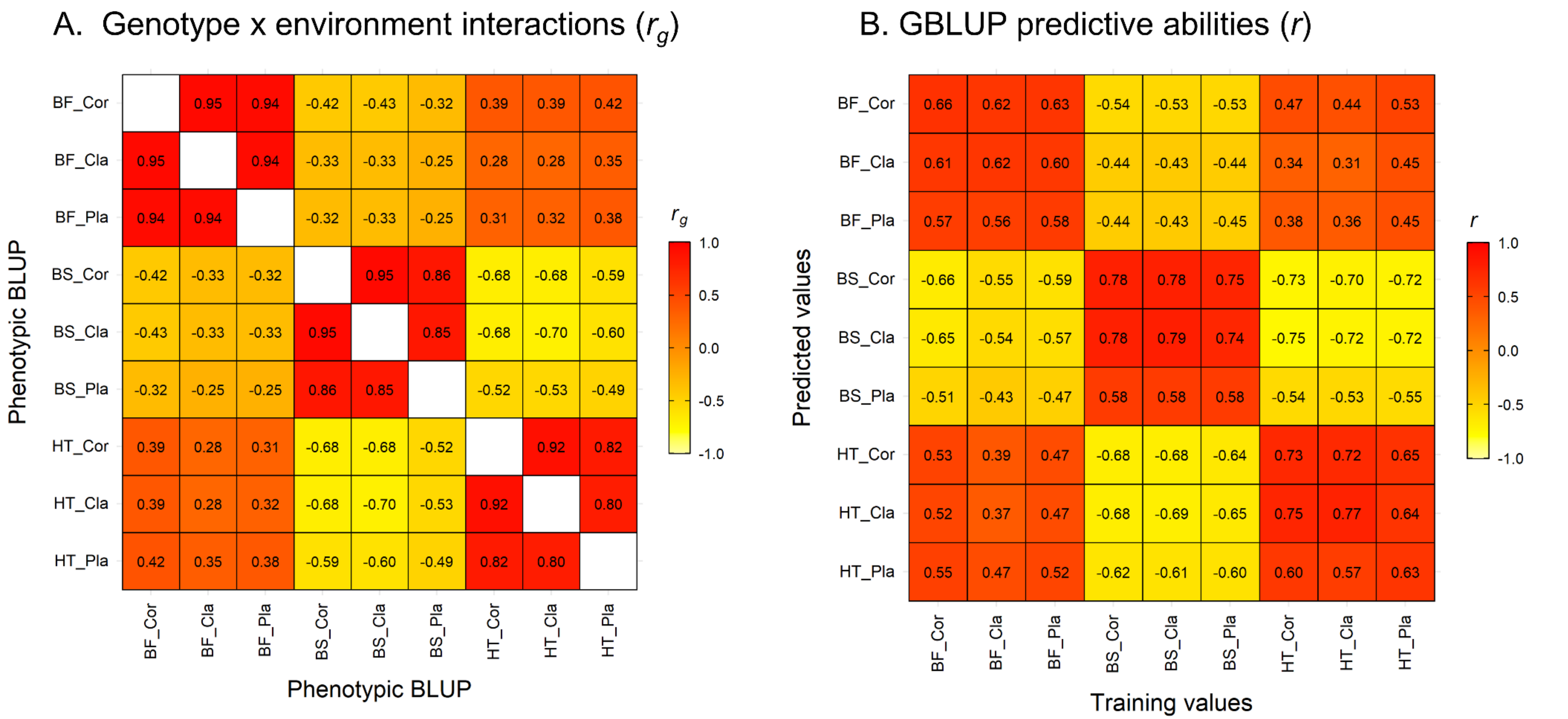


**Figure S5. Extent of genotype by environmental (GxE) interactions based on among-plantation genetic correlations and predictive abilities.** (A) Genetic correlations among bud flush (BF), bud set (BS), and height growth (HT) within (main diagonal) and across (off-diagonal values) the Corvallis (Cor), Clatskanie (Cla), and Placerville (Pla) test plantations. (B) Corresponding GBLUP predictive abilities across all hierarchical levels (genotypes, stands, and rivers) based on 20,770,783 single-nucleotide polymorphism markers filtered using 'liberal' criteria (Table S4). All predictive abilities are averages from 100 random, ten-fold cross-validations with a training population size of 756 and prediction population size of 84.

### A. Vegetative bud flush

| Stage | Description | Examples |
|---|---|---|
| 0 | Buds tightly closed | 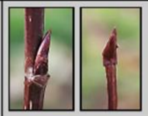 |
| 1 | Buds swollen/breaking/leaf primordia visible (green visible between scales) | 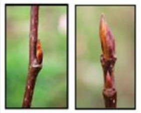 |
| 2 | Leaves emerging (any leaves showing outside of bud scales) | 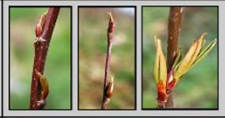 |
| 3 | Leaves fully emerged and unfolding (leaf petiole is visible) | 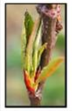 |
| 4 | Leaves fully unfolded | 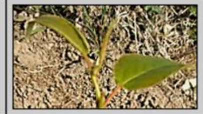 |
| 5 | Leaves fully expanded | 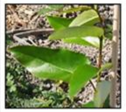 |

### B. Vegetative bud set

| Stage | Description | Examples |
|---|---|---|
| 1 | Actively growing<br>Many large new leaves at the apex | 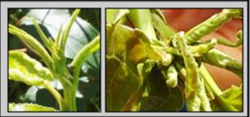 |
| 2 | Slowing down<br>Some new leaves at the apex | 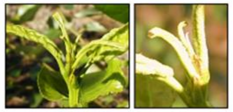 |
| 3 | Just prior to bud set<br>One small new leaf at the apex<br>Enlarged stipules | 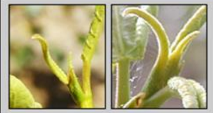 |
| 4 | Just after bud set<br>Stipules form a point at the apex with no new leaves protruding | 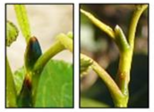 |
| 5 | Small reddish bud | 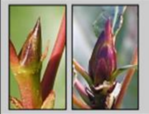 |
| 6 | Large reddish and woody bud | 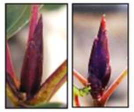 |

**Figure S6. Scores used to quantify the timing of vegetative bud flush and bud set in *Populus trichocarpa*.** In this species, the bud scales are formed from modified stipules.

### Supporting Materials and Methods

*Plant materials and test plantations*

We assembled a study population of 1101 black cottonwood clonal genotypes by combining existing and new collections. First, we obtained 282 genotypes collected from western Oregon and western Washington by GreenWood Resources, Inc. (Portland, OR) and 215 genotypes collected from southwestern British Columbia by the British Columbia Ministry of Forests. We also collected new cuttings from large, continuous stands of black cottonwood along four rivers in northwestern Oregon and western Washington. We collected genotypes along the Columbia ($N$ = 140), Puyallup ($N$ = 161), Skykomish ($N$ = 141), and Skagit ($N$ = 140) Rivers, and from five small, isolated stands near Lake Tahoe in California ($N$ = 22). Collections were designed to sample populations that had a common demographic history and were not affected by recent introgression from other *Populus* species (i.e., Figs. 1-1 and 1-2 in DIFAZIO *et al.* (2011). We focused on the portion of the range where black cottonwood has optimal growth, high levels of phenotypic variation, and weak interpopulation differentiation for neutral markers (WEBER AND STETTLER 1981; WEBER *et al.* 1985; DEBELL 1990; SLAVOV *et al.* 2012).

Cuttings collected from the field or arboreta were rooted individually in 115-ml plastic Cone-tainers (SC7, Stuewe & Sons, Inc., Tangent, OR) under greenhouse conditions (Mt. Jefferson Farms, Inc., Salem, OR). Rooted cuttings were overwintered in an outdoor shadehouse at Oregon State University, before outplanting at three sites. The northernmost plantation was at a site owned by GreenWood Resources near Clatskanie, OR. Another plantation was established near Oregon State University in Corvallis. The southernmost test plantation was established at the USDA Forest Service Nursery near Placerville, CA. At each plantation, a single copy of each genotype was planted in each of three randomized complete blocks, and the plantations were fenced to prevent deer and livestock damage. Trees were planted at 3 × 3 m spacing at the Clatskanie and Placerville plantations, and 2 × 3 m at the Corvallis plantation. Because of the tighter spacing at the Corvallis plantation, it was coppiced after the 2010 growing season. Irrigation was not needed at the Clatskanie plantation, the Corvallis plantation was overhead-irrigated biweekly during the first growing season, and the Placerville plantation (the driest site) was drip-irrigated weekly during each growing season. Plants that died during the first growing season were replanted in the summer of 2010 with filler trees, but these trees were not used in the statistical analyses.

*Phenotypic data analysis*

Because the blocks were large (i.e., up to ~1100 trees), we accounted for microsite variation using spatial analysis, rather than using a block term in the statistical model. To achieve homogeneity of variance among plantations, we then standardized the raw data at each plantation using the phenotypic standard deviation (WHITE 1996). For single-site analyses, we used these adjusted data and a hierarchical random effects model to calculate variance components and random effects (i.e., best linear unbiased predictors, BLUPs). This model included terms for the plantation mean, river (R), stand-within-river [S(R)], genotype-within-stand-and-river [G(SR)], and error. The across-site model included additional terms for the random effect of plantation and interactions of plantation with R, S(R), and G(SR). These analyses involved five steps. First, for each plantation, we adjusted the raw data using SAS PROC TPSPLINE (SAS v9.4, SAS Institute Inc., Cary, NC). Second, we used a statistical model with genotype as a random effect to estimate phenotypic variance ($\sigma^2_{\mathrm{p}}$) as $\sigma^2_{\mathrm{G}} + \sigma^2_{\varepsilon}$, where $\sigma^2_{\mathrm{G}}$ was the genotype variance component and $\sigma^2_{\varepsilon}$ was the error variance estimated using SAS PROC GLIMMIX. Third, to achieve homogeneity of variance among plantations, we then standardized the raw data at each plantation using the phenotypic standard deviation (WHITE 1996). Fourth, we adjusted the standardized raw data, again using PROC TPSPLINE. Finally, for the single-site analyses, we used these adjusted data and the following statistical model to calculate variance components and random effects [best linear unbiased predictors (BLUPs)] using PROC GLIMMIX.

$$X_{\mathrm{jklm}} = \mu + R_{\mathrm{j}} + S_{\mathrm{k}}(R_{\mathrm{j}}) + G_{\mathrm{l}}(SR_{\mathrm{kj}}) + \varepsilon_{\mathrm{jklm}}, \quad (1)$$

where $X_{jklm}$ is the observation for the m$^{th}$ tree of the l$^{th}$ genotype from the k$^{th}$ stand within the j$^{th}$ river; $\mu$ is the overall mean; $R_j$ is the random effect of the j$^{th}$ river; $S_k(R_j)$ is the random effect of the k$^{th}$ stand within the j$^{th}$ river; $G_l(SR_{kj})$ is the random effect of the l$^{th}$ genotype from the k$^{th}$ stand and j$^{th}$ river; and $\varepsilon_{jklm}$ is the experimental error. However, before running the final analysis, we used this same model to identify and remove outliers that had residuals exceeding three standard deviations.

We used the results from Model 1 primarily to ascertain data quality, whereas all remaining analyses were conducted using the results from the following across-site model:

$$X_{ijklm} = \mu + P_i + R_j + S_k(R_j) + G_l(SR_{kj}) + P_i \cdot R_j + P_i \cdot S_k(R_j) + P_i \cdot G_l(SR_{kj}) + \varepsilon_{ijklm}, \quad (2)$$

where $X_{ijklm}$ is the observation for the m$^{th}$ tree of the l$^{th}$ genotype from the k$^{th}$ stand within the j$^{th}$ river at the i$^{th}$ plantation; $P_i$ is the random effect of the i$^{th}$ plantation; $P_i \cdot R_j$, $P_i \cdot S_k(R_j)$, and $P_i \cdot G_l(SR_{kj})$ are the interactions between the i$^{th}$ plantation and the j$^{th}$ river, k$^{th}$ stand within the j$^{th}$ river, and the l$^{th}$ genotype from the k$^{th}$ stand within the j$^{th}$ river; and other effects are as described above.

For each *P. trichocarpa* genotype, we calculated two "phenotypes," $G(SR)$ and $G$, by combining random effects from Model 2. $G(SR)$ includes only the $G(SR)$ random effect, whereas $G$ includes the random effects from all three hierarchical levels, $G(SR)$ and $S(R)$, and $R$. We then estimated genetic correlations among traits (within and among plantations) as the Pearson product-moment correlations among these values.

We used the variance components from Model 2 to calculate heritabilities and characterize the distribution of genetic variation at different spatial scales. We estimated variance components for rivers ($\sigma^2_{\mathrm{R}}$), stands-within-rivers ($\sigma^2_{\mathrm{S(R)}}$), genotypes-within-stands-and-rivers ($\sigma^2_{\mathrm{G(SR)}}$), and error ($\sigma^2_{\varepsilon}$). $\sigma^2_{\mathrm{G(SR)}}$ represents within-stand genetic variation, whereas the sum of $\sigma^2_{\mathrm{R}}$, $\sigma^2_{\mathrm{S(R)}}$, and $\sigma^2_{\mathrm{G(SR)}}$ is the total genetic variation ($\sigma^2_{\mathrm{G}}$). Additionally, we estimated variance components for the plantation (environment) by genetic interactions, $\sigma^2_{\mathrm{R \cdot E}}$, $\sigma^2_{\mathrm{S(R) \cdot E}}$, and $\sigma^2_{\mathrm{G(SR) \cdot E}}$, the sum of which is the total genotype-by-environmental (GxE) variance ($\sigma^2_{\mathrm{G \cdot E}}$).

We calculated total heritabilities as $H_G^2 = \sigma_{\mathrm{G}}^2/(\sigma_{\mathrm{G}}^2 + \sigma_{\mathrm{G\cdot E}}^2 + \sigma_{\varepsilon}^2)$, and within-stand heritabilities as $H_{G(SR)}^2 = \sigma_{\mathrm{G(SR)}}^2/(\sigma_{\mathrm{G(SR)}}^2 + \sigma_{\mathrm{G(SR)\cdot E}}^2 + \sigma_{\varepsilon}^2)$. Proportions of genotypic variation were calculated as $\sigma_{\mathrm{R}}^2/\sigma_{\mathrm{G}}^2$ for rivers, $\sigma_{\mathrm{S(R)}}^2/\sigma_{\mathrm{G}}^2$ for stands-within-rivers, and $\sigma_{\mathrm{G(SR)}}^2/\sigma_{\mathrm{G}}^2$ for genotypes-within-stands. To further characterize genetic structure, we calculated among-stand $Q_{\mathrm{ST}}$ values at each hierarchical level. $Q_{\mathrm{ST}}$, a measure of allelic differentiation for quantitative traits, was calculated as follows (WHITLOCK 2008; WHITLOCK AND GILBERT 2012): $Q_{\mathrm{ST}} = (\sigma_{\mathrm{R}}^2 + \sigma_{\mathrm{S(R)}}^2)/(\sigma_{\mathrm{R}}^2 + \sigma_{\mathrm{S(R)}}^2 + 2\sigma_{\mathrm{G(SR)}}^2)$. Finally, to illustrate spatial patterns of genetic structure, we performed principal component analysis (PCA) of the data matrix consisting of $G$ values for the three phenotypic traits in each of the three test plantations (i.e., nine variables). This was done using the *prcomp* function in R (TEAM 2019), with the *center* and *scale* options set to TRUE.

*SNP data*

To remove related trees, we first clustered the initial set of 970 individuals into seven groups using fastSTRUCTURE (RAJ *et al.* 2014). Next, we calculated a genomic relationship matrix (GRM) for each group using the *--make-grm-gz* option of GCTA (YANG *et al.* 2011). Finally, we removed one individual from each pair that had a GRM value exceeding 0.25. We excluded 42 other individuals for four reasons. We removed 12 individuals because they could not be grouped into stands and rivers with the other clonal genotypes (note, 'river' denotes a river or drainage). Second, we removed all 13 individuals from an atypical, strongly differentiated subpopulation from Lake Tahoe (pairwise $F_{\mathrm{ST}} > 0.15$) (SLAVOV *et al.* 2012; EVANS *et al.* 2014). Third, we removed all individuals from three rivers that had < 10 individuals each (Kitimat, Nisqually, and Cowlitz), resulting in the removal of another 14 individuals. Finally, we removed three individuals that did not cluster within their respective rivers based on the first two SNP principal components (PCs) (i.e., were > 3 standard deviations from the river centroids). The remaining 840 clonal genotypes represented 91 stands in 16 rivers (Fig. 1).

For the final analyses, we filtered SNPs based on three sets of criteria (Table S4) using VCFtools v. 0.1.14 (DANECEK *et al.* 2011) and PLINK v.1.90b4.4 (CHANG *et al.* 2015). We used 'strict' filtering to set SNP genotypes (AA, AB, BB) with < 8 reads or genotype quality < 20 as missing data (*vcftools --minDP* 8 *--minGQ* 20), and then excluded SNPs with minor allele frequency (MAF) < 0.01 (*vcftools --maf* 0.01) or > 10% missing data (*plink --geno* 0.1). For maximum genome coverage, we used 'liberal' filtering to retain all SNPs with at least 2 copies of the minor allele (*vcftools --mac* 2), and did not filter based on the number of reads per genotype, genotype quality, proportion of missing data, or MAF. Finally, we also simulated a set of 51,820 'RAD-Seq' SNPs based on high-quality RAD loci that were detected in each of seven *P. alba* and seven *P. tremula* genotypes using *PstI* and a minimum sequencing coverage of 6× (STOELTING *et al.* 2013). We translated the positions of these known loci from version 2 to version 3 of the *P. trichocarpa* genome (TUSKAN *et al.* 2006), and then selected all bi-allelic, non-singleton SNPs from our re-sequencing data that were within 100-bp of a *PstI* restriction site. Because *PstI* is methylation sensitive, some loci are not 'RAD-accessible,' and will be missed. Thus, this approach masks our re-sequencing data to simulate the RAD-accessible portion of the genome.

*Linkage disequilibrium and haplotype sharing*

We calculated $r^2$ for each pair of SNPs located within 10 kb of each other using PLINK (*plink --r2 --ld-window 1000 --ld-window-kb 10 --ld-window-r2 0*) using different MAF cut-offs or bins. The MAF bins (Table S5) were set so that all pairs of loci in a bin had potentially high LD [i.e., an $r^2$ with a theoretical maximum of at least 0.5 (WRAY 2005)]. To estimate the probability of tagging a QTN, we first selected $M = 10^2 – 10^7$ SNPs at random using PLINK (*plink --thin-count*), then randomly selected a putative QTN from the $M$ SNPs and calculated $r^2$ between the QTN and all SNPs located within 10 kb from it. This was repeated 100 times for each value of $M$ and the estimate of tagging probability was calculated as the proportion of times at least one SNP within 10 kb had $r^2 \geq 0.6$ with the putative QTN.

To compare LD within and among rivers, we randomly sampled 10 clonal genotypes from each river and 1 genotype from each of 10 randomly sampled rivers. This approach equalized the within-river and among-river comparisons (i.e., the minimum sample size within river was $N$ = 10), which is important because $r^2$ is sensitive to sample size (TENESA *et al.* 2007). Next, we calculated pairwise $r^2$ as described above using 10 replications. For these analyses, we eliminated singleton SNPs from each sample (MAF ≥ 0.10), calculated $r^2$ only for SNPs that were separated by at least 300 bp, and then corrected $r^2$ for the small sample size by subtracting 1/2$N$.

To quantify linkage phase consistency, we first selected a subset of SNPs with MAF ≥ 0.01 that were separated by at least 300 bp and located on chromosomes 1-19. We then calculated the genotypic correlations ($g_r$) between all pairs of SNPs from that set that were located within 10 kb of each other using PLINK (*plink --r --ld-window 50 --ld-window-kb 10*). To calculate haplotype sharing among rivers ($hs_R$), we calculated $g_r$ values for each river, calculated the correlation between $g_r$ values for each pair of rivers ($r_{gr}$), and then calculated $hs_R$ as the overall average of all $r_{gr}$ values across all pairs of rivers. We used an analogous approach to calculate haplotype sharing among stands-in-rivers ($\underline{hs_{S(R)}}$), but only for the Columbia, Skykomish, and Skagit rivers. These were the only rivers that had multiple stands with at least 20 clones each. Finally, we calculated haplotype sharing among genotypes-within-stands-and-rivers ($\underline{hs_{G(SR)}}$) by sampling four stands (each in a different river) that had at least 40 clones and randomly splitting clones in each stand into two groups of at least 20 clones to calculate within strand $r_{gr}$. To minimize the effects of allele frequency differentiation on haplotype sharing, we also repeated this analysis for a set of SNPs with 0.01 ≤ MAF < 0.11 in each of the 16 rivers. Because very few SNPs satisfied these criteria ($M$ = 1002), we extended the window in which we calculated haplotype sharing to 1 Mb. Standard errors of haplotype sharing ($hs$) means were calculated as $[var(hs)/k]^{1/2}$, where $k$ is the number of pairs of rivers, stands or samples of genotypes within stands for which $hs$ values were calculated.

*Random forest and ridge regression analyses*

We used random forest regression and ridge regression to compare the abilities of climatic and geographic variables to predict quantitative traits and SNP PC scores at the stand (S) and river (R) levels. For each approach, we calculated predictive abilities for four sets of predictor variables: (1) geographic variables (latitude, longitude, and elevation), (2) 21 temperature and precipitation related variables from ClimateNA v5.21 (1961-1990 normals; WANG *et al.* 2016), (3) a subset of climate variables selected using random forest regression at the stand level, and (4) all geographic and climatic variables.

To select the most important climate variables (i.e., climate variable subset), we first used random forest regression at the stand level. These analyses resulted in a separate set of selected climate variables for each phenotypic trait and SNP PC score (Table S1). Next, we used these selected climate variables and the other predictors described above to calculate PA as the correlation between the phenotypic BLUP or SNP PC score versus the predictions from the regression models. After the independent variables were centered and scaled using the R *scale* function, both linear and squared values were included in the models. Random forest regression was conducted using the *randomForest* function from the *randomForest* R package v. 4.6.14 (LIAW AND WIENER 2002) using default parameters, ntree = 2000, sampsize = 2/3, and replace = FALSE. Ridge regression was conducted using the *cv.glmnet* function from the *glmnet* R package (FRIEDMAN *et al.* 2010) using default parameters, s = lambda.min, and alpha = 0 (ridge regression). These functions were run using a random cross-validation approach with 8 folds at the river level ($N$ = 16) and 10 folds at the stand level ($N$ = 91). Each cross-validation analysis was run 100 times, and the mean correlation (PA) and its standard error were calculated.

*Delineation of seed zones*

Zones were delineated and reconstructed by clustering the stand-level PC scores of phenotypes, SNPs, climate variables, and geographic variables using the R *kmeans* function, with the *centers* option set to the number of desired zones and *nstart* set to 25. To compare the true vs reconstructed zone allocations, we calculated cluster purity (MANNING *et al.* 2008), which is the proportion of stands in each reconstructed seed zone that were also in the same true zone. The phenotypic data consisted of stand-level averages for the first 5 PCs derived from a PCA of 9 phenotypes (i.e., 3 traits measured in 3 plantations). SNP data consisted of stand-level averages for the first 5 PCs derived from a PCA of the GRM calculated using SNPs filtered with the 'liberal' criteria (Table S4). Climate and geographic data consisted of stand-level averages for the first 5 PCs (climate) or 3 PCs (geography) derived from PCAs of the same variables used for the random forest and ridge regression analyses descried above (*SI Dataset* S1)

**References**


Chang, C. C., C. C. Chow, L. C. Tellier, S. Vattikuti, S. M. Purcell *et al.*, 2015 Second-generation PLINK: rising to the challenge of larger and richer datasets. Gigascience 4**:** 7.

Danecek, P., A. Auton, G. Abecasis, C. A. Albers, E. Banks *et al.*, 2011 The variant call format and VCFtools. Bioinformatics 27**:** 2156-2158.

DeBell, D. S., 1990 Populus trichocarpa Torr. & Gray, Black Cottonwood, pp. 570-576 in *Silvics of North America Vol. 2. Hardwoods. Agriculture Handbook 654*, edited by R. M. Burns and B. H. Honkala. U.S. Department of Agriculture, Forest Service, Washington D.C.

DiFazio, S. P., G. T. Slavov and C. P. Joshi, 2011 Populus : A premier pioneer system for plant genomics pp. 1-28 in *Genetics, Genomics and Breeding of Poplar* edited by C. P. Joshi, S. P. DiFazio and C. Kole. Science Publishers, Enfield, NH.

Evans, L. M., G. T. Slavov, E. Rodgers-Melnick, J. Martin, P. Ranjan *et al.*, 2014 Population genomics of Populus trichocarpa identifies signatures of selection and adaptive trait associations. Nat Genet 46**:** 1089-1096.

Friedman, J., T. Hastie and R. Tibshirani, 2010 Regularization Paths for Generalized Linear Models via Coordinate Descent. J Stat Softw 33**:** 1-22.

Liaw, A., and M. Wiener, 2002 Classification and Regression by randomFores, pp. 18-22 in *R News*, *https://CRAN.R-project.org/doc/Rnews/*.

Manning, D. C., P. Raghavan and H. Schütze, 2008 *Introduction to Information Retrieval*. Cambridge University Press, Cambridge, UK.

Raj, A., M. Stephens and J. K. Pritchard, 2014 fastSTRUCTURE: Variational Inference of Population Structure in Large SNP Data Sets. Genetics 197**:** 573.

Slavov, G. T., S. P. DiFazio, J. Martin, W. Schackwitz, W. Muchero *et al.*, 2012 Genome resequencing reveals multiscale geographic structure and extensive linkage disequilibrium in the forest tree Populus trichocarpa. New Phytologist 196**:** 713-725.
Stoelting, K. N., R. Nipper, D. Lindtke, C. Caseys, S. Waeber *et al.*, 2013 Genomic scan for single nucleotide polymorphisms reveals patterns of divergence and gene flow between ecologically divergent species. Molecular Ecology 22**:** 842-855.
Team, R. C., 2019 R: A language and environment for statistical computing, pp. R Foundation for Statistical Computing, URL http://www.R-project.org/, Vienna, Austria.
Tenesa, A., P. Navarro, B. J. Hayes, D. L. Duffy, G. M. Clarke *et al.*, 2007 Recent human effective population size estimated from linkage disequilibrium. Genome Research 17**:** 520-526.
Tuskan, G. A., S. DiFazio, S. Jansson, J. Bohlmann, I. Grigoriev *et al.*, 2006 The Genome of Black Cottonwood, Populus trichocarpa (Torr. & Gray). Science 313**:** 1596-1604.
Wang, T., A. Hamann, D. Spittlehouse and C. Carroll, 2016 Locally Downscaled and Spatially Customizable Climate Data for Historical and Future Periods for North America. PLOS ONE 11**:** e0156720.
Weber, J. C., and R. F. Stettler, 1981 Isoenzyme variation among ten [riparian] populations of Populus trichocarpa Torr. et Gray in the Pacific Northwest. Silvae Genetica 30**:** 82-87.
Weber, J. C., R. F. Stettler and P. E. Heilman, 1985 Genetic variation and productivity of Populus trichocarpa and its hybrids. I. Morphology and phenology of 50 native clones. Canadian Journal of Forest Research 15**:** 376-383.
White, T., 1996 Genetic parameter estimates and breeding value predictions: issues and implications in tree improvement programs, pp. 110-117 in *Tree improvement for sustainable tropical forestry QFRI IUFRO Conference,* edited by M. J. Dieters, A. C. Matheson, D. G. Nikles, C. E. Harwood and S. M. Walker, Caloundra, Queensland, Australia.
Whitlock, M. C., 2008 Evolutionary inference from Q(ST). Molecular Ecology 17**:** 1885-1896.
Whitlock, M. C., and K. J. Gilbert, 2012 QST in a hierarchically structured population. Molecular Ecology Resources 12**:** 481-483.
Wray, N. R., 2005 Allele frequencies and the r2 measure of linkage disequilibrium: impact on design and interpretation of association studies. Twin Res Hum Genet 8**:** 87-94.
Yang, J., S. H. Lee, M. E. Goddard and P. M. Visscher, 2011 GCTA: A Tool for Genome-wide Complex Trait Analysis. American Journal of Human Genetics 88**:** 76-82.
